# Structure of the pre-mRNA leakage 39-kDa protein reveals a single domain of integrated zf-C3HC and Rsm1 modules

**DOI:** 10.1101/2022.08.11.503642

**Authors:** Hideharu Hashimoto, Daniel H. Ramirez, Ophélie Lautier, Natalie Pawlak, Günter Blobel, Benoît Palancade, Erik W. Debler

## Abstract

In *Saccharomyces cerevisiae*, the pre-mRNA leakage 39-kDa protein (*Sc*Pml39) was reported to retain unspliced pre-mRNA prior to export through nuclear pore complexes (NPCs). Pml39 homologs outside the *Saccharomycetaceae* family are currently unknown, and mechanistic insight into Pml39 function is lacking. Here we determined the crystal structure of *Sc*Pml39 at 2.5 Å resolution to facilitate the discovery of orthologs beyond *Saccharomycetaceae*, e.g. in *Schizosaccharomyces pombe* or human. The crystal structure revealed integrated zf-C3HC and Rsm1 modules, which are tightly associated through a hydrophobic interface to form a single domain. Both zf-C3HC and Rsm1 modules belong to the Zn-containing BIR (Baculovirus IAP repeat)-like super family, with key residues of the canonical BIR domain being conserved. Features unique to the Pml39 modules refer to the spacing between the Zn-coordinating residues, giving rise to a substantially tilted helix aC in the zf-C3HC and Rsm1 modules, and an extra helix αAB’ in the Rsm1 module. Conservation of key residues responsible for its distinct features identifies *S. pombe* Rsm1 and *Homo sapiens* NIPA/ZC3HC1 as structural orthologs of *Sc*Pml39. Based on the recent functional characterization of NIPA/ZC3HC1 as a scaffold protein that stabilizes the nuclear basket of the NPC, our data suggest an analogous function of *Sc*Pml39 in *S. cerevisiae*.

## Introduction

Nuclear pore complexes (NPCs) are large macromolecular assemblies (MW of ~60 MDa in the budding yeast *Saccharomyces cerevisiae*) and mediate the transport of a tremendous range of cargoes such as water, ions, small molecules, proteins, and ribonucleoparticles across the nuclear envelope^1–4^. As exclusive transport channels between the nucleus and cytoplasm, NPCs are ideally positioned to serve as gateways or checkpoints in the flow of information from DNA to protein and have therefore long been postulated to function in “gene gating”^5^. NPCs indeed play key roles beyond nucleocytoplasmic transport, such as in genome organization and integrity, gene regulation, and mRNA quality control (QC)^6–12^.

The mRNA QC ensures that incompletely processed and/or improperly assembled mRNA ribonucleoparticles (mRNPs) are discarded to avoid detrimental effects on protein homeostasis or RNA metabolism^13,14^. This processing status and export competency of mRNPs can be monitored through their association with the nuclear basket, which is envisioned to serve as a platform to which mRNPs transiently associate prior to nuclear export to the cytoplasm^11,15–19^. For example, the poly(A)-binding protein *Sc*Nab2, which directly binds to the C-terminal region of nuclear basket of the NPC protein Myosin-like protein 1 (*Sc*Mlp1), monitors proper 3’-mRNA processing^15,20–22^. In turn, *Sc*Mlp1 deletion triggers cytoplasmic leakage of intron-containing pre-mRNAs^23^. A similar role in the nuclear retention of unspliced reporter mRNAs *in vivo* was assigned to homologs in fission yeast (*Sp*Nup211) and mammals (Tpr)^24–26^. Furthermore, a pre-mRNA leakage phenotype was observed for the deletion of an *Sc*Mlp1/*Sc*Mlp2-interacting protein termed *S. cerevisiae* pre-mRNA leakage protein 39-kDa (*Sc*Pml39)^27^. Conversely, its overexpression traps intron-containing mRNAs in nuclear foci enriched in *Sc*Mlp1 and *Sc*Nab2^27^. *Sc*Pml39 is required for cell growth in the absence of a functional Y-complex, an essential NPC building block required for mRNA export^28,29^, and is recruited to the nuclear basket of NPCs by virtue of interactions with the N-terminal regions of *Sc*Mlp1 and *Sc*Mlp2^27^. While *Sc*Mlp1/2 and *Sc*Nab2 sequences and functions are highly conserved across phyla^11,21,30^, the *Sc*Pml39 sequence is unique to *Saccharomycetaceae* (**Fig. S1**). To date, no homologous sequences could be found in human, *S. pombe*, and others. Thus, identifying orthologs in organisms beyond *Saccharomycetacea* would accelerate our understanding of the function of *Sc*Pml39.

As form follows function, determining the atomic structure of *Sc*Pml39 and identifying key residues involved in its structural integrity is a course of action to find *Sc*Pml39 orthologs^31^. The structure prediction programs Phyre2 and SWISS-MODEL^32,33^ identified two <u>B</u>aculoviral <u>I</u>nhibitor of apoptosis (IAP) <u>R</u>epeat (BIR) domains in *Sc*Pml39. BIR domains contribute to protein-protein interactions in both apoptotic^34–38^ and non-apoptotic pathways^39–41^. The canonical BIR domain is well-studied, with currently 312 structures available in the Protein Data Bank (PDB), including structures in complex with substrate peptides. Their structures are highly conserved, as illustrated by a root-mean square deviation (RMSD) of Ca atoms of less than 1.0 Å for the representative BIR domains human neuronal apoptosis inhibitor protein (PDB: 2VM5), human Survivin (PDB: 3UED), and *D. melanogaster* IAP-BIR1 (**Fig. S2a**)^42^. The BIR domain is comprised of ~70 amino acid residues, three a-helices (αA, αB, and αC) and one β-sheet formed by three anti-parallel strands (β1, β2, and β3). The BIR domain harbors a cysteine-cysteine-histidine-cysteine (CCHC)-type zinc finger (ZnF) motif with the consensus sequence Rxx(S/T)Ω…GΩ…**C**-x_2_-**C**-x_16_-**H**-x_6_-**C**-x-Ω (x denotes any amino acid residue and Ω an aromatic residue) (**Fig. S2a**). The importance of the conserved residues for structural integrity was previously noted^43^. In particular, the motif Rxx(S/T)Ω is located in helix αA, and the Arg residue is essential. The motif GΩ is located between αB and β1 to form a sharp β-turn. The first and second Zn-coordinating cysteine are in the loop between β1 and β2, the Zn-coordinating histidine is in helix αC, and the last Zn-coordinating cysteine is in the loop between helices αC and αD. The final aromatic residue Ω is located in helix αE to undergo π-cation interactions with Arg in the Rxx(S/T)Ω motif of helix αA (**Fig. S2a**).

In *Sc*Pml39, the two potential ZnF motifs deviate from the consensus sequence by a shortened linker between the Zn-coordinating histidine and cysteine residues: C-x_2_-C-x_n_-H-**x_3_**-C. This difference is a hallmark of the zf-C3HC (ID: PF07967) and Rsm1 (ID: PF08600) protein families, which together with the canonical BIR domain (ID: PF00653) comprise the CCHC ZnF motif-containing clan of BIR-like domains (ID: CL0417) according to the Pfam database^44^. *Sc*Pml39 is predicted to have two zfC3HC and/or Rsm1 domains. In contrast to the canonical BIR and zf-C3HC domains, the Rxx(S/T)Ω motif could not be identified in the Rsm1 family by a Hidden Markov model (HMM)^45^ (**Fig. S3**). Furthermore, altough the zfC3HC and Rsm1 domains are widely distributed in 1897 sequences from 1208 species and 1155 sequences from 896 species in the Pfam database, respectively, a structure of these domains is currently not available in the PDB. Since the spacing between the ZnF-coordinating residues is critical for ZnF structure and function^46^, we set out to determine the crystal structure of *Sc*Pml39 to assess the impact of the ZnF motif differences on the zf-C3HC and Rsm1 domains.

Here we present the 2.5 Å-resolution crystal structure of *Sc*Pml39 and identify orthologs in *S. pombe* and human based on structure-guided sequence alignment. The distinct spacing between the CCHC ZnF-coordinating residues results in features that are different from the canonical BIR domain. Two zf-C3HC and Rsm1 modules tightly associate to form a novel “Pml39 fold”.

## Results

### ScPml39 contains a single domain that recapitulates subcellular localization, function, and ScMlp1-interaction of full-length ScPml39

The domain structure of *Sc*Pml39 was determined by limited proteolysis using the full-length recombinant protein *Sc*Pml39 (residues 1-334). Elastase digest yielded a stable single fragment of ~30 kDa molecular weight (**Fig. 1a**). Based on the domain boundaries identified by mass spectrometry, we generated a truncated *Sc*Pml39 construct comprising residues 77-317 for structural studies. Notably, *Sc*Pml39_77-317_ is a functional domain fragment both *in vitro* and *in vivo*. In yeast cells, GFP-tagged *Sc*Pml39_77-317_ expressed in *pml39*Δ mutant yeast cells was recruited to the NPC nuclear basket and displayed the typical U-shaped perinuclear staining as the wild type (**Fig. 1b,c**)^27^. Expression of *Sc*Pml39_77-317_-GFP and full-length *Sc*Pml39-GFP similarly complemented the synthetic growth defect arising from the simultaneous loss-of-function of *Sc*Pml39 and of the Y-complex (*nup133*Δ) (**Fig. 1d**). *in vitro, Sc*Pml39_77-317_ maintains direct binding to a recombinant homodimer of N-terminal fragment of *Sc*Mlp1 (residues 1-325) with a dissociation constant (*K*_D_) of ~13 μM and a 1:1 molar ratio, as measured by isothermal titration calorimetry (**Fig. 1e,f**).

**Figure 1.**
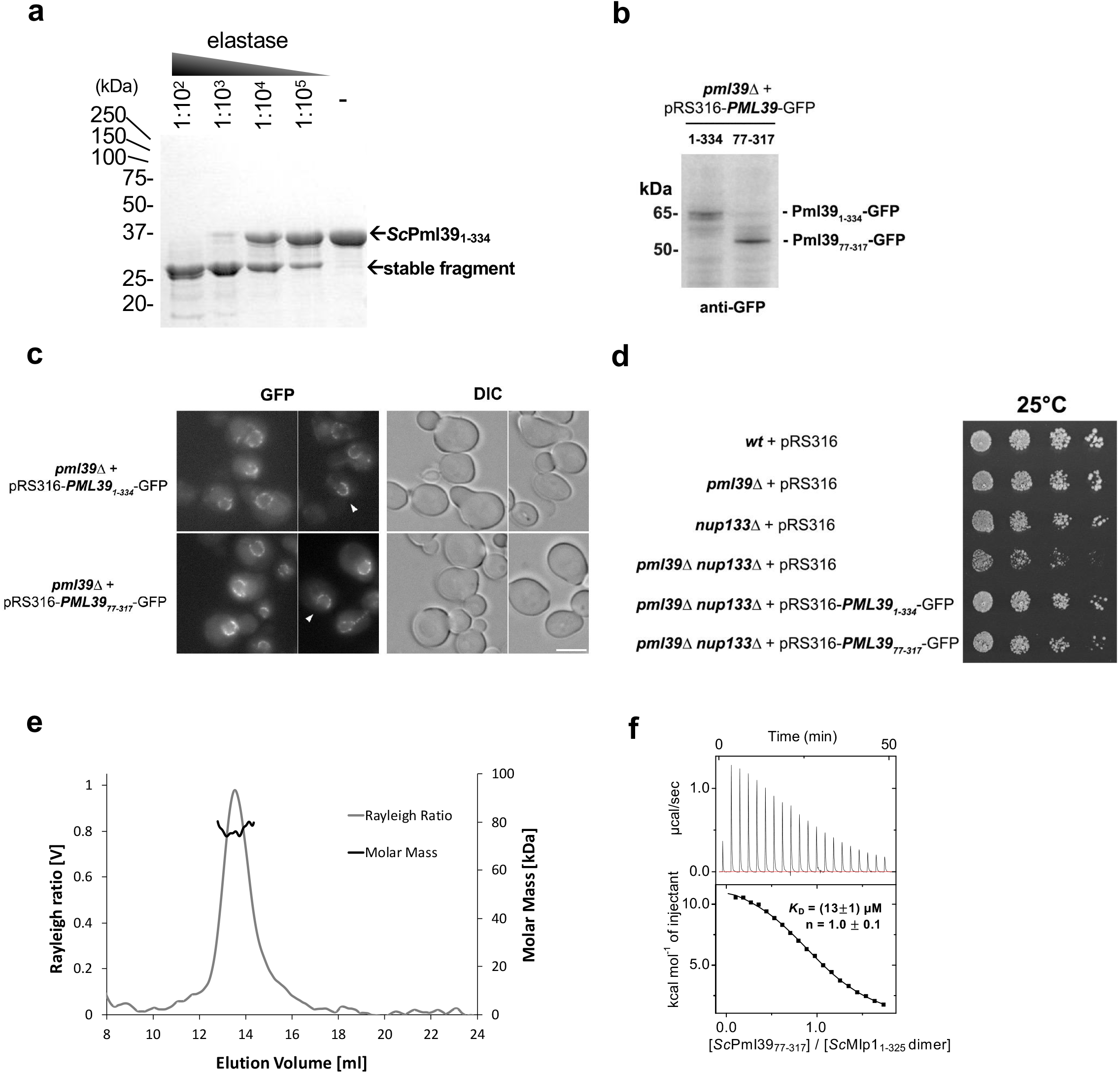
*Sc*Pml39 contains a single domain that recapitulates subcellular localization, function, and *Sc*Mlp1-interaction of full-length *Sc*Pml39. **a)** Limited proteolysis of recombinant full-length *Sc*Pml39 at the indicated dilutions of 2 mg/ml elastase analyzed by SDS-PAGE. The full-length version of the cropped gel is presented in **Fig. S4a**. **b)** Expression of GFP-tagged full-length (1-334) or truncated (77-317) versions of *Sc*Pml39 in *pml39*Δ cells detected by immunoblotting using anti-GFP antibodies. The full-length version of the cropped blot is presented in **Fig. S4b**. **c)** Live imaging of *pml39*Δ cells expressing GFP-tagged full-length (1-334) or truncated (77-317) versions of *Sc*Pml39. Single plane images are shown for the GFP and DIC (differential interference contrast) channels. Arrowheads point to nuclei showing the U-shaped perinuclear staining typical of *Sc*Pml39. Scale bar, 5 μm. **d)** Cells of the indicated genotypes were spotted as serial dilutions on SC medium and grown for 3 days at 25°C. **e)** Recombinant *Sc*Mlp1_1-325_ forms a homodimer. Size exclusion chromatography coupled to multi-angle light scattering (SEC-MALS) using a Superdex 200 10/300 column was used. Molecular mass determination and Rayleigh ratio of *Sc*Mlp1_1-325_ (light and dark gray, respectively) demonstrated that the *Sc*Mlp1_1-325_ (expected molecular size is 41 kDa) has a dimeric size (~80 kDa). **f)** Isothermal titration calorimetry (ITC) thermogram (upper panel) and plot of corrected heat values (lower panel) showed that monomeric *Sc*Pml39_77-317_ binds dimeric *Sc*Mlp1_1-325_ at a 1:1 molar ratio with a *K*_D_ value of ~13 μM.

### ΔN-ScPml39 contains two tightly interacting BIR-like Rsm1 and zf-C3HC modules

*Sc*Pml39_77-317_ was crystalized in the *P*3_1_21 space group. The phases and the structure were determined to a resolution of 2.5 Å by Zinc single-wavelength anomalous dispersion (Zn-SAD) (**Table 1** and **Fig. S5**). The crystallographic asymmetric unit contains one molecule. The majority of the fragment (residues 79-311, Fig. 2a) was well resolved in the electron density, while no electron density was observed for two N-terminal residues (77-78), residues 148-151 between β3 strand and helix αC, a Ser-rich region comprising residues 213-226, and six C-terminal residues (312-317, **Fig. 2b**). *Sc*Pml39_77-317_ contains two BIR-like modules: zf-C3HC (residues 79-189 in blue) and Rsm1 (residues 190-311 in purple) (**Fig. 2b,c**). The two zf-C3HC and Rsm1 modules form a single structural domain stabilized by the hydrophobic residues Leu87, Ile90, Pro111, Leu112, Leu185, Tyr189 of zf-C3HC and Phe272 and Trp291 of Rsm1 (**Fig. 3**). Each module could not be expressed individually as a soluble protein in *E. coli* (data not shown), consistent with the limited proteolysis data (**Fig. 1a**).

**Table 1.** Data collection and refinement statistics.

|  |  |
| --- | --- |
| **Data collection** |  |
| Beamline | 8.2.2 (ALS) |
| Space group | P3<sub>1</sub>21 |
| Cell dimensions |  |
| a, b, c (Å) | a=53.0, b=53.0, c=171.3 |
| $\alpha$, $\beta$, $\gamma$ (°) | $\alpha=\beta=90$, $\gamma=120$ |
| Wavelength (Å) | 1.282 |
| Resolution (Å) <sup>a</sup> | 44.34-2.49 (2.58-2.49) |
| No. of unique reflections | 18,841 (1,837) |
| R<sub>merge</sub> (%) <sup>a,b</sup> | 4.5 (88.9) |
| CC1/2 <sup>a</sup> | 1 (0.954) |
| CC* <sup>a</sup> | 1 (0.988) |
| $\langle I / \sigma I \rangle$ <sup>c</sup> | 30.4 (3.2) |
| Completeness (%) <sup>a</sup> | 99.6 (99.7) |
| Redundancy <sup>b</sup> | 11.3 (11.4) |
| **Refinement** |  |
| Resolution (Å) | 44.34 – 2.49 |
| No. of reflections | 18,812 |
| Test set | 941 |
| R<sub>work</sub> <sup>d</sup> / R<sub>free</sub> <sup>e</sup> (%) | 23.3 / 26.3 |
| No. of atoms |  |
| Protein | 1779 |
| Zn | 2 |
| R.m.s. deviations |  |
| Bond lengths (Å) | 0.003 |
| Bond angles (°) | 0.49 |
| <B-Value> (Å<sup>2</sup>) |  |
| Protein | 97.0 |
| Zn | 80.6 |
| Ramachandran plot <sup>f</sup> |  |
| Favored (%) | 98.1 |
| Allowed (%) | 1.9 |
| Outliers (%) | 0.0 |
| Rotamer outliers (%) | 0 |
| Clashscore | 4.2 |
| Cβ deviation | 0 |
| PDB | 7RDN |

<sup>a</sup>Highest-resolution shell is shown in parentheses.
<sup>b</sup>R<sub>merge</sub> = $\sum |I - \langle I \rangle| / \sum I$ , where $I$ is the observed intensity and $\langle I \rangle$ is the averaged intensity from multiple observations.
<sup>c</sup> $\langle I / \sigma I \rangle$ = averaged ratio of the intensity ( $I$ ) to the error of the intensity ( $\sigma I$ ).
<sup>d</sup>R<sub>work</sub> = $\sum |F_{obs} - F_{cal}| / \sum |F_{obs}|$ , where $F_{obs}$ and $F_{cal}$ are the observed and calculated structure factors, respectively.
<sup>e</sup>R<sub>free</sub> was calculated using a randomly chosen subset (5%) of the reflections not used in refinement.
<sup>f</sup>As determined by MolProbity.

**Figure 2.**
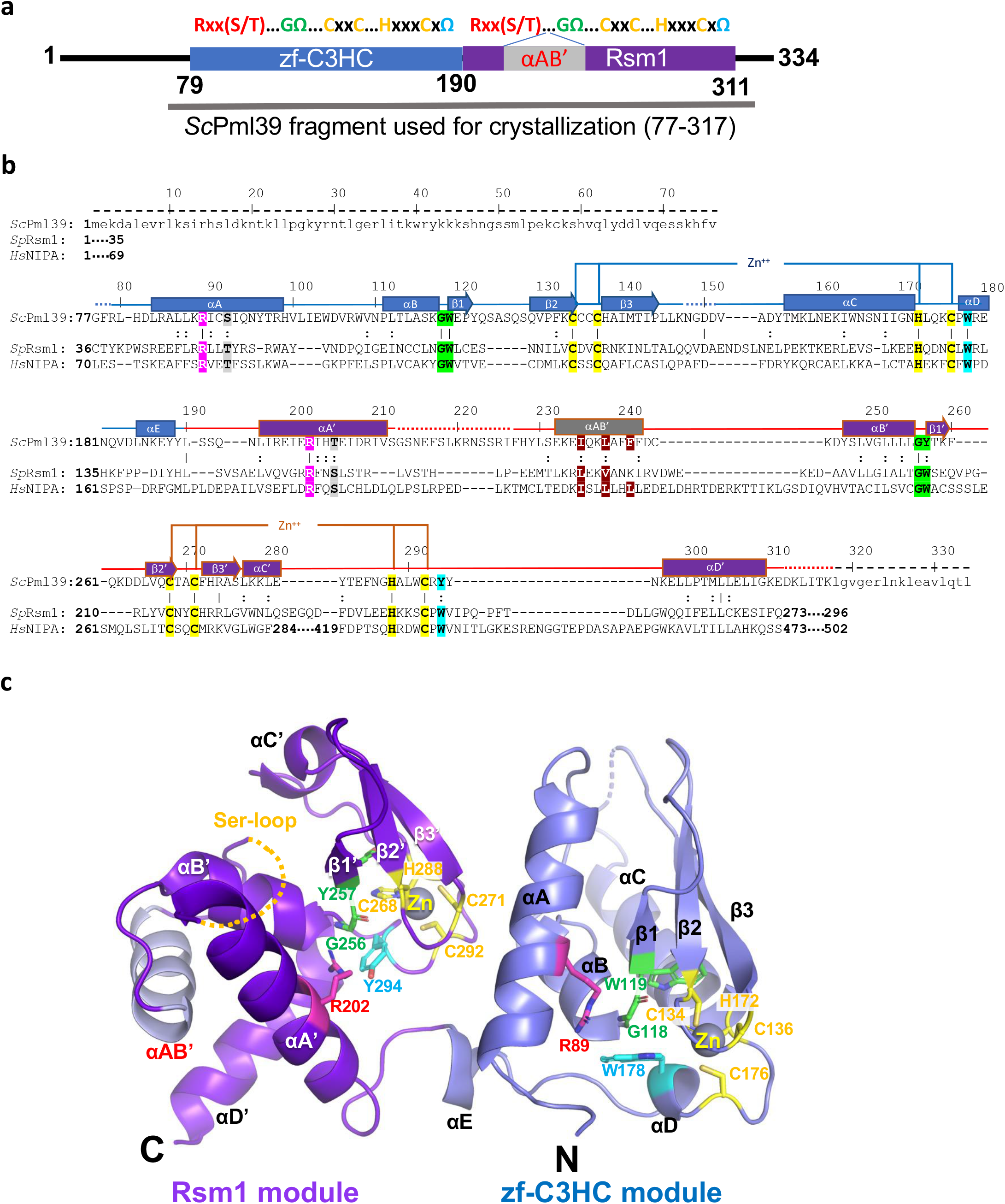
Crystal structure of *Sc*Pml39_77-317_. **a)** Schematic of *Sc*Pml39. The consensus sequence of the canonical BIR domain is shown on the top. The zf-C3HC and Rsm1 modules are indicated in blue and purple, respectively. The fragment used for crystallization (residues 77-317) is shown. **b)** Structure-guided sequence alignment of *Sc*Pml39, *Sp*Rsm1 and human NIPA/ZFC3HC1. αA–αE refer to α-helices, and β1–β3 to β-strands, indicating the secondary structure elements of *Sc*Pml39. Residue numbering is shown for *Sc*Pml39. Residues highlighted designate conserved Arg (magenta) and Ser/Thr (gray) in αA, Gly-aromatic residues between αB and β1 (green), conserved zinc-coordinating residues (yellow), and a conserved aromatic residue in αE (cyan). Similar and identical residues are marked as : and |, respectively. Disordered regions are represented by dotted lines, whereas regions lacking in the crystallization fragment are indicated by dashed lines and lowercase letters. **c)** The Pml39 fold (cartoon representation) and side chains of key residues (stick representation) are shown.

**Figure 3.**
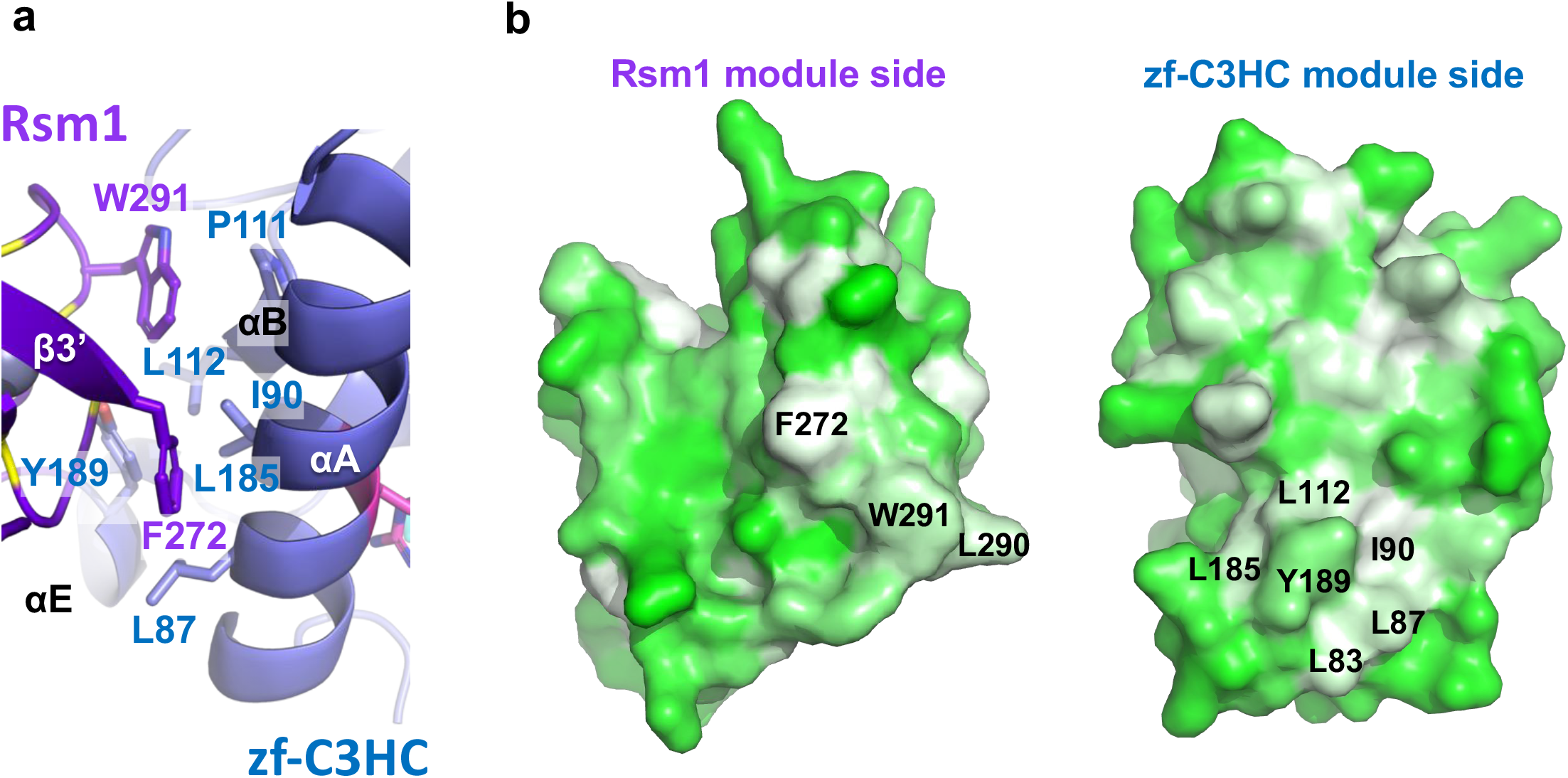
Hydrophobic interface between zf-C3HC and Rsm1 modules. **a)** Residues involved in van der Waals’ contacts are shown in stick representation. **b)** Hydrophobicity of Rsm1 and zf-C3HC module are shown using Pymol script color_h, ranging from white (highly hydrophobic) to green (less hydrophobic). Residues indicated in a) are shown.

### *Structure of the* zf-C3HC *module of ScPml39*

The *Sc*Pml39 zf-C3HC module comprises five a-helices (αA (83-98), αB (111-117), αC (156-171), αD (177-180), and αE (185-189)) and one antiparallel β-sheet composed of three strands (β1 (119-121), β2 (129-134), and β3 (138-144)) (**Figs. 2, S6**). Arg89 and Ser92 in helix αA are part of the conserved Rxx(S/T)Ω consensus motif. The guanidinium group of Arg89 interacts with the aromatic ring of Trp178 in helix αD, as observed in canonical BIR domain proteins (**Fig. 4b**). A 12-residue loop (His99 to Asn110) connects the helices αA and αB. Gly118 forms a sharp turn and connects helix αB and strand β1. The adjacent aromatic residue Trp119 is part of an invariant GΩ dipeptide motif present in the canonical BIR domains^36,43,47^. Trp119 and Met114 in the strand β3 engage in a sulphur-aromatic interaction, and an internal hydrophobic network within the β-sheet, helices αB, αD, and αC stabilize the zf-C3HC module (**Fig. 4c)**. While these characteristics classify the zf-C3HC domain as a member of the BIR-like family, the CCHC ZnF motif is different both in sequence and in structure from the canonical BIR domain (**Fig. S2**). Cys134, Cys137, His172, and Cys176 residues coordinate Zn to form the C-x_2_-C-x-H-**x_3_**-C ZnF motif (**Fig. 2c**). The three residues between His172 and Cys176 represent a unique signature of zf-C3HC domain proteins, in contrast to six residues in canonical BIR domain proteins (**Figs. S1, S2**). Furthermore, the presence of 34 residues between Cys137 and His172 in Pml39 zf-C3HC module, compared to 16 residues in canonical BIR domains, results in an elongated zf-C3HC module comprising ~110 residues versus ~70 residues in most canonical BIR domains^36,48,49^.

**Figure 4.**
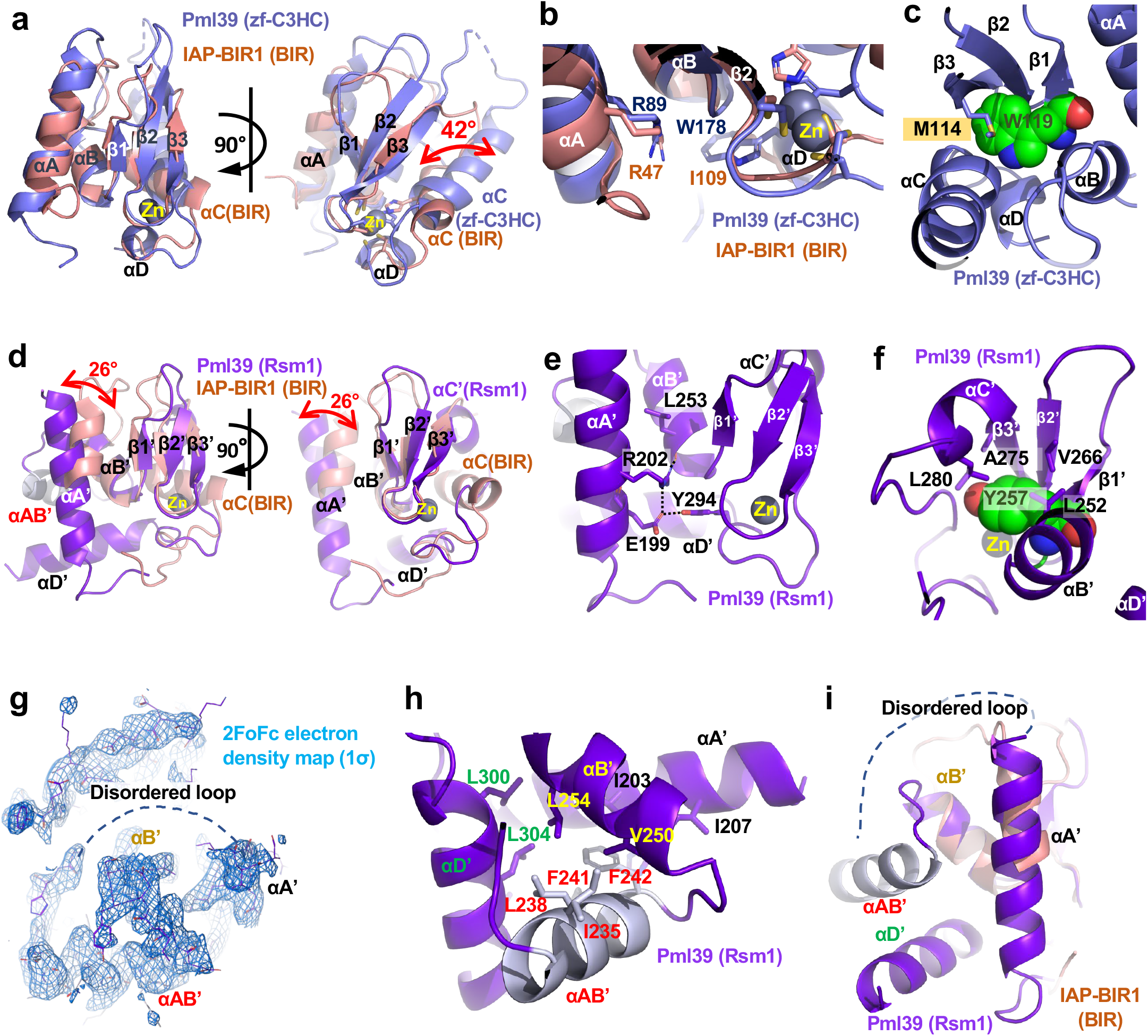
Structure of *Sc*Pml39 modules. **a)** Superimposition of *Sc*Pml39 zf-C3HC module (in blue) and a representative canonical BIR domain, *D. melanogaster* IAP-BIR1 domain structure (in orange, PDB: 3SIP) (left panel) and a view rotated by ~90° (right panel). **b)** Structural conservation of Arg89 in helix αA and Trp178 in helix αD. **c)** Internal hydrophobic network in *Sc*Pml39 zf-C3HC module. **d)** Superimposition of *Sc*Pml39 Rsm1 module (in purple) and a representative canonical BIR domain, *D. melanogaster* IAP-BIR1 domain structure (in orange, PDB: 3SIP) (left panel) and a view rotated by ~90° (right panel). **e)** Structural conservation of Arg202 in helix αA’ and Tyr294 in helix αD’. **f)** Internal hydrophobic network in *Sc*Pml39 Rsm1 module. **g)** 2FoFc electron density, contoured at 1σ above the mean for helix αAB’ in Rsm1 module. **h)** Internal hydrophobic network in *Sc*Pml39 Rsm1 module helix αAB’. **i)** Different view from h).

The Pml39 zf-C3HC module superimposes with *D. melanogaster* IAP-BIR1 (PDB: 3SIP), a representative structure of a canonical BIR domain, with an RMSD of 2.4 Å (**Fig. 4a**). The Zn ion of the Pml39 zf-C3HC module adopts almost the same position as in canonical BIR domain proteins. Due to the different spacing within the CCHC ZnF motif, helix αC of the *Sc*Pml39 zf-C3HC module is tilted by 42° with respect to the corresponding helix αC in *D. melanogaster* IAP-BIR1 domain, rendering the distinct spacing a key determinant for the topology of *Sc*Pml39 (**Fig. 4a**). Finally, the conserved aromatic residue succeeding the CCHC ZnF motif – Trp178 in the zf-C3HC module – packs against the C-terminal end of helix αB like a lid and forms numerous hydrophobic contacts in the core of the module (**Figs. 2c, 4b**).

### Structure of the Rsm1 module of ScPml39

The Rsm1 module consists of 122 residues and comprises five a-helices (αA’ (196-211), αAB’ (232-242), αB’ (247-255), αC’ (277-281), and αD’ (297-310)) and one antiparallel β-sheet composed of three β-strands (β1’ (257-259), β2’ (265-268), and β3’ (272-276). The topology of the *Sc*Pml39 Rsm1 module deviates even more from canonical BIR domains than the *Sc*Pml39 zf-C3HC module (**Figs. 2c, S6**). The additional helix αAB’ between the helices αA’ and αB’ extends the sequence of the Rsm1 module and alters its topology (**Figs. 4d, S6**).

Sequence analysis alone could not detect the N-terminal BIR consensus sequence motif Rxx(S/T)Ω (**Fig. S3**), but the structure of the *Sc*Pml39 Rsm1 module indeed reveals the presence of this motif in helix αA’, containing Arg202 and Thr205 (**Fig. 2**). In addition to the conserved hydrogen bond of the Arg202 guanidinium group with the carboxylate Oη atom of Glu199, the guanidinium group forms a hydrogen bond with the main-chain oxygen of Leu253 instead of the hydroxyl group of Tyr294 (**Fig. 4e**). Thus, this residue plays a similar role for the structural integrity of the Rsm1 module as the corresponding arginine in canonical BIR domain proteins.

The extensive insertion between helices αA’ and αB’ is unique to the Rsm1 module among the three BIR-like domain families. While the Ser-rich region immediately preceding helix αAB’ is disordered, the electron density of helix αAB’ is clearly observed, and its C-terminus is connected to helix αB’ through a short segment (Asp243-Tyr247) (**Figs. 2b, 4g-i**). Ile236, Leu239, Phe241, and Phe242 in helix αAB’ tether and stabilize helices αA’, αB’, and αE’ through hydrophobic interactions (**Fig. 4h**). The insertion of helix αAB’ tilts helix αA’ by 26° with respect to the corresponding helix in canonical BIR domains, such as *D. melanogaster* IAP-BIR1 (**Fig. 4d)**. The region between helix αB’ and strand β3’ of the Rsm1 module aligns well with the corresponding region of IAP-BIR1 and contains the highly conserved GΩ motif (Gly256-Tyr257) between helix αB’ and strand β1’. Leu252 of helix αB’, Tyr257 of strand β1’, Val266 of strand β2’, Ala275 of strand β3’, and L280 of helix αC’ form hydrophobic interactions that stabilize the domain structure (**Fig. 4f**).

Cys268, Cys271, His288, and Cys292 coordinate Zn and are part of the C-x_2_-C-x**_16_**-H-x**_3_**-C ZnF motif (**Figs. 2b, S1**). The Zn ion of the Rsm1 module adopts the same position as the Zn ion in canonical BIR domains. Similar to the zf-C3HC module, the distinct CCHC ZnF spacing impacts the structure of the Rsm1 module, with helix αC’ being displaced with respect to canonical BIR domains (**Fig. 4d**). Thus, the distinct CCHC ZnF spacing between Zn-coordinating residues and helix αAB’ insertion between helices αA’ and αB’ represent key determinants of the Rsm1 module. The conserved aromatic residue succeeding the CCHC ZnF motif – Tyr294 in the Rsm1 module – packs against the C-terminal end of helix αB’ like a lid and contributes numerous hydrophobic interactions in the core of the module (**Figs. 2c, 4e**).

### Schizosaccharomyces pombe Rsm1 and human NIPA/ZC3HC1 are structural orthologs of ScPml39

*Sc*Pml39 harbors an architecture of two consecutive zf-C3HC and Rsm1 modules. Using structure-guided sequence analysis, we identified *S. pombe* Rsm1 (UniProtKB: O94506)^50,51^ and *H. sapiens* nuclear-interacting partner of ALK (*Hs*NIPA/ZC3HC1, UniProtKB: Q86WB0)^38,52,53^ as *Sc*Pml39 structural orthologs. Both *Sp*Rsm1 and *Hs*NIPA/ZC3HC1 feature a domain organization with two tandem zf-C3HC/Rsm1 modules and meet the criteria for conservation of *Sc*Pml39 residues essential for structural integrity (**Fig. 2b**). In *Sp*Rsm1, Arg49 and Thr52 would correspond to the *Sc*Pml39 residues Arg89 and Ser92 in helix αA, respectively, Gly74-Trp75 of *Sp*Rsm1 would correspond to the Gly118-Trp119 motif, Cys85, Cys88, His126, and Cys130 in *Sp*Rsm1 would coordinate the zinc ion, and the conserved Trp132 would correspond to Trp178 of *Sc*Pml39. Arg156 and Ser159 would be in the helix αA’ of the Rsm1 module. The helix αAB’ seems difficult to predict, but the extended sequence of *Sp*Rsm1 in this region is shared with *Sc*Pml39. Gly204-Trp205 would locate between helix αB’ and strand β2’. Cys216, Cys219, His241, and Cys245 would coordinate zinc, followed by Trp247. Thus, *Sp*Rsm1 harbors all conserved residues related to the structural integrity of *Sc*Pml39. zf-C3HC and Rsm1 modules can also be identified in the human NIPA/ZC3HC1 sequence, which can be aligned with that of *Sc*Pml39. As for the zf-C3HC module, Arg81 and Thr84 would be in helix αA, Gly106-Trp107 would connect helix αB and strand β1, Cys118, Cys120, His152, and Cys156 would coordinate the zinc ion, and Trp158 would be the final conserved aromatic residue of the consensus sequence in the zf-C3HC module. As for the Rsm1 module, Arg185 and Ser188 would be in helix αA’, Gly255-Trp256 would connect helix αB’ and strand β1’, Cys272, Cys275, His425, and Cys459 would coordinate the zinc ion, and an aromatic residue (Trp461) following Cys459 is also conserved in its putative Rsm1 module. Therefore, the NIPA/ZC3HC1 structure is expected to be highly similar to that of *Sc*Pml39 (**Fig. 2c**), except for an extensive (~135-residue) region that is inserted between the Zn-coordinating Cys275 and His425. This analysis suggests that the Pml39 structure is not unique to *Saccharomycetaceae*, but also found in *S. pombe* and in humans.

## Discussion

The *Sc*Pml39_77-317_ crystal structure revealed two zf-C3HC and Rsm1 modules that tightly interact to form a single domain termed “Pml39 fold”. Our analysis suggests that the Pml39 fold is not an architecture unique to *Saccharomycetaceae*, but is likely to exist in 934 proteins whose sequences contain tandem zf-C3HC and Rsm1 modules across all phyla in the Pfam database^44^.

While all three families of the BIR-like clan (canonical BIR, zf-C3HC, and Rsm1 domains) share conserved key residues responsible for the structural domain integrity, the ZnF motif in the zf-C3HC and Rsm1 families with the consensus sequence C-x_2_-C-x_n_-H-x_3_-C is distinct from the canonical BIR domain (C-x_2_-C-x_n_-H-x_6_-C). Moreover, the additional helix αAB’ insertion is solely found in the Rsm1 module. These findings together with the difficulty to identify homologs on the sequence level suggest that the Pml39 fold has rapidly evolved, as only key residues are conserved. A possible sequence of events for the evolution of the Pml39 fold is as follows: 1) mutation in the CCHC zinc finger motif of an ancestral BIR domain, 2) domain duplication, 3) insertion of helix αAB’ and extra residues between helix αC and the zinc-coordinating histidine in the Rsm1 module. Steps of insertion/deletion are expected to make it more difficult for sequence algorithms to predict the Pml39 fold^53^. Indeed, the N-terminal consensus motif of the Rsm1 module, Rxx(S/T)Ω, could not be clearly identified *in silico* (**Fig. S3**).

The structure of *Sc*Pml39 has enabled us to unambiguously identify *S. pombe Sp*Rsm1 and human NIPA/ZC3HC1 as structural orthologs of *Sc*Pml39 (**Fig. 2b**). AlphaFold2 also predicts the Pml39 fold for *Sp*Rsm1 and human NIPA/ZC3HC1 (**Fig. S7)**^54^. Strikingly, the overall amino acid sequence identity/similarity is very low, with only key residues being conserved among *Sc*Pml39, *Sp*Rsm1, and human NIPA/ZC3HC1 (**Fig. 2b**). As it is not known whether the function is conserved among these proteins as well, we tested if *Sp*Rsm1 could rescue the *Sc*Pml39-deficient yeast cell phenotype. Under this heterologous condition, GFP-tagged *Sp*Rsm1 did not localize to the nuclear periphery (**Fig. S8a)**. In addition, *Sp*Rsm1 expression does not complement the *nup133Δ / pml39*Δ synthetic interaction in the growth assay (**Fig. S8b**)^27,55^. Low sequence identity/similarity in the *Sc*Pml39 interacting N-terminal region of Mlp1 (**Fig. 1f**) may be a barrier to compensate *Sc*Pml39 function by *Sp*Rsm1 expression in the budding yeast cells, providing a possible reason for the failure of the rescue assay (**Fig. S9**). Indeed, genetic studies support our hypothesis that *Sp*Rsm1 is involved in mRNA export^50,51^. Moreover, human NIPA/ZC3HC1 has been identified as a nuclear basket-associated protein, required to scaffold Tpr polypeptides^56,57^. These data, together with the identification of *Sp*Rsm1 and human NIPA/ZC3HC1 as structural *Sc*Pml39 orthologs, suggest a function of *Sc*Pml39 as a scaffold protein to stabilize the nuclear basket. We conclude that *Sc*Pml39 is likely conserved in structure and function from yeast to vertebrates.

## Materials and Methods

### Protein expression and purification

DNA fragments encoding full-length (residues 1-334) and truncated (residues 77-317) *Saccharomyces cerevisiae* Pml39 (UniProtKB: Q03760) and a DNA fragment encoding a C-terminally truncated (residues 1-325) construct of *Saccharomyces cerevisiae* Mlp1 (UniProtKB: Q02455) were amplified by PCR from genomic DNA and cloned into the *NcoI/NotI* restriction sites of the pET28a vector (Novagen). The constructs were overexpressed in *E. coli* BL21(DE3)-RIL CodonPlus cells (Agilent Technologies) and grown in LB media containing appropriate antibiotics. Protein expression was induced at O*D*_600_ of ~0.6 with 0.1 mM isopropyl-β-D-thiogalactoside (IPTG) at 18° C for 16 h. The cells were harvested by centrifugation at 7,500 x *g* and 4 °C and lysed with a cell disruptor (Avestin) in a buffer containing 20 mM Tris-HCl, pH 8.0, 300 mM NaCl, 14.3 mM β-mercaptoethanol, 0.5 mM 4-(2-aminoethyl)benzenesulfonyl fluoride hydrochloride (AEBSF) (Sigma), 2 μM bovine lung aprotinin (Sigma), and complete EDTA-free protease inhibitor cocktail (Roche). After centrifugation at 35,000 x *g* for 45 min, the cleared lysate was loaded onto a Ni-NTA column (Qiagen) and eluted with an imidazole gradient. Protein-containing fractions were pooled, dialyzed against a buffer containing 20 mM Tris-HCl, pH 8.0, 5 mM dithiothreitol (DTT), and 250 mM NaCl for full-length *Sc*Pml39, 100 mM NaCl for *Sc*Pml39_77-317_, or 150 mM NaCl for *Sc*Mlp1_1-325_, and subjected to cleavage with PreScission protease (GE Healthcare) for 5 h at 4° C. Following hexahistidine-tag removal, *Sc*Pml39 proteins were bound to a HiTrap SP column (GE Healthcare) and eluted with a NaCl gradient. For ΔC-*Sc*Mlp1, a HiTrap Q column was used. Protein-containing fractions were pooled, concentrated, and purified on a HiLoad Superdex 200 (16/60) gel filtration column (GE Healthcare) in a buffer containing 20 mM HEPES-NaOH, pH 7.5, 200 mM NaCl, and 1 mM Tris(2-carboxyethyl)phosphine hydrochloride (TCEP). Protein concentrations was measured by absorbance at 280 nm, protein was flash-frozen in liquid nitrogen and stored at −80 °C.

### Limited proteolysis

In a volume of 100 μl, full-length *Sc*Pml39 at 1.3 mg/ml was incubated with a dilution series of 2 mg/ml porcine elastase at room temperature for 30 min. An aliquot of each dilution was mixed with reducing SDS-PAGE sample buffer and analyzed by SDS-PAGE. The remaining reaction volumes were quenched by guanidinium chloride powder for mass spectrometry analysis. To this end, the samples were run over a reversed phase column (PLRP-S), collected peaks were injected into an ion trap mass spectrometer, and spectra were analyzed by GPMAW^58^.

### Crystallization, data collection, structure determination, and refinement

Crystals of *Sc*Pml39_77-317_ were grown at 20° C in hanging drops containing 1 μL of the protein at 10 mg/ml and 1 μL of a reservoir solution consisting of 12% (w/v) PEG 8,000 and 0.1 M HEPES-NaOH, pH 7.7. Crystals grew in space group *P*3_1_21 within a week, were cryo-protected in 25% (v/v) glycerol containing 12% (w/v) PEG 8,000, and 0.1 M HEPES-NaOH, pH 7.7, and flash-cooled in liquid nitrogen. X-ray diffraction data were collected at the beamlines X29A at the National Synchrotron Light Source (NSLS) of the Brookhaven National Laboratory (BNL) and 8.2.2 at the Advanced Light Source (ALS). Diffraction data were processed in HKL2000^59^. The structure was solved by the single anomalous dispersion (SAD) phasing technique running the script AutoSol of the PHENIX package^60^. The asymmetric unit contained one molecule. Model building was performed in O^61^ and Coot^62^. The final model spanning residues 79-310 was refined in PHENIX to *R*_free_ / *R*_work_ factors of 26.3% / 23.3% with excellent stereochemistry and clash score as assessed by MolProbity^63^. Details for data collection and refinement statistics are summarized in Table 1. Figures were generated using PyMOL (Schrödinger, LLC), the electrostatic potential was calculated with APBS^64^. Atomic coordinates and structure factors have been deposited with the Protein Data Bank under PDB: 7RDN.

### Isothermal titration calorimetry

ITC measurements were performed at 4 °C using a MicroCal auto-iTC200 calorimeter (GE Healthcare). Samples were extensively dialyzed against a buffer containing 500 mM NaCl, 20 mM HEPES (pH 7.5), and 0.5 mM TCEP. After dialysis, the protein was filtered (0.22 μm) and centrifuged, followed by determining their concentration by UV absorbance at 280 nm. 2 μL of 1.6 mM *Sc*Pml39_77-317_ was injected into 350 μL of 70 μM *Sc*Mlp1_1-325_ in the chamber every 180 s. Baseline-corrected data were analyzed using the ORIGIN software to determine the molar ratio (n), dissociation constant (*K*_D_), and enthalpy (ΔH). These parameters were subsequently used to determine the free Gibbs energy (ΔG) and the entropic component (TΔS) using ΔG = -RT ln (1/*K*_D_) and TΔS = ΔH – ΔG equations, where R and T are the gas constant (1.99 cal/(mol*K)) and absolute temperature, respectively. Thermodynamic parameters are represented as mean values ± standard deviation calculated from three independent measurements.

### Size-Exclusion Chromatography and Multi-Angle Light Scattering (SEC-MALS)

SEC experiments were performed on 100 μL injections of 70 μM *Sc*Mlp1_1-325_ with a Superdex 200 (10/300) GL column (GE Healthcare) at 0.5 mL min^−1^ at 25°C in 500 mM NaCl, 20 mM HEPES-NaOH (pH 7.5), and 0.5 mM TCEP. Absolute molecular weights were determined using MALS. The scattered light intensity of the column eluant was recorded at 18 different angles using a DAWN-HELEOS MALS detector (Wyatt Technology Corp.) operating at 658 nm after calibration with the monomer fraction of Type V BSA (Sigma). Protein concentration of the eluant was determined using an in-line Optilab T-rex interferometric refractometer (Wyatt Technology Corp.). The weight-averaged molecular weight of species within defined chromatographic peaks was calculated using the ASTRA software version 6.0 (Wyatt Technology Corp.), by construction of Debye plots (KC/*R*θ *versus* sin^2^[θ/2]) at 1-s data intervals. The weight-averaged molecular weight was then calculated at each point of the chromatographic trace from the Debye plot intercept, and an overall average molecular weight was calculated by averaging across the peak.

### Yeast strains and plasmids

All *S. cerevisiae* strains used in this study (Table S1) are haploid, isogenic to BY4742 and were obtained by transformation and/or successive crosses using standard procedures. pRS316-*PML39_1-334_-GFP*, pRS316-*PML39_77-317_-GFP* and pRS316-*RSM1-GFP* expression vectors were constructed by PCR-based techniques using *Saccharomyces cerevisiae* or *Schizosaccharomyces pombe* genomic DNA and pFA6a-GFP(S65T)-KanMX as templates^65^. Expression of the three transgenes is driven by the *PML39* endogenous promoter (300 bp upstream the ATG codon). Unless indicated, cells were grown at 30 °C in rich (YPD, Yeast Extract Peptone Dextrose) or Synthetic Complete (SC) media^27,55^ and harvested during exponential phase.

### *in vivo* assays

Localization of tagged fluorescent proteins was analyzed in live cells grown in SC medium. Wide-field fluorescence images of GFP-tagged versions of *Sc*Pml39 or *Sp*Rsm1 were acquired using a Leica DM6000B microscope with a 100X/1.4 NA (HCX Plan-Apo) oil immersion objective and a CCD camera (CoolSNAP HQ; Photometrics), and further scaled equivalently using the MetaMorph software (Molecular Devices). Whole-cell extracts were prepared from cells grown in YPD and analyzed by SDS-PAGE using stain-free precast gels (Biorad) followed by westernblotting with monoclonal anti-GFP antibodies (clones 7.1 & 13.1, Sigma)^66^. Growth assays were achieved by spotting serial dilutions of cells on SC medium and incubating the plates at 25° C.

### Data Availability

The datasets and materials used and/or analyzed during the current study available from the corresponding author on reasonable request. Atomic coordinates and structure factors have been deposited with the Protein Data Bank under PDB: 7RDN.

## Acknowledgements

We thank David King (University of California, Berkeley) for mass spectrometry analysis, Corrie Ralston and Peter Zwart for support during data collection at the Advanced Light Source (ALS), Wuxian Shi for support during data collection at of the National Synchrotron Light Source (NSLS), Kushol Gupta from the Johnson Foundation Structural Biology and Biophysics Core at the Perelman School of Medicine (Philadelphia, PA) for performing the SEC-MALS analysis, Anna Mienko for assistance with protein expression and purification, the High-Throughput Screening and Spectroscopy Resource Center at Rockefeller University, and the X-Ray Crystallography & Molecular Interactions Facility at the Sidney Kimmel Cancer Center, which is supported in part by National Cancer Institute Cancer Center Support Grant P30 CA56036 and S10 OD017987. X-ray data were collected at beamline 8.2.2 of the ALS, a U.S. DOE Office of Science User Facility under Contract No. DE-AC02-05CH11231, supported in part by the ALS-ENABLE program funded by the National Institutes of Health, National Institute of General Medical Sciences, grant P30 GM124169-01, and at beamline X29 of NSLS. This work was supported by funds from the Rockefeller University and the Howard Hughes Medical Institute (to G.B.), by a grant from the Agence Nationale pour la Recherche (ANR-18-CE12-0003, to B.P.), and by a National Institute of Allergy and Infectious Diseases (NIAID) grant from the National Institutes of Health (NIH) (R01AI165840, to E.W.D.). We would like to thank the members of the Debler laboratory for helpful discussions.

## Author contributions

E.W.D. and B.P. designed the research, interpreted the data, and edited the manuscript. H.H. and E.W.D. performed the x-ray crystal analysis. N.P., D.H.R., and E.W.D. performed protein expression, protein purification, and crystallization. D.H.R. and E.W.D. performed ITC measurements. O.L. and B.P. performed *in vivo* experiments. E.W.D., B.P., and G.B. provided funding. G.B. provided valuable feedback in the early stages of the project. H.H. drafted the manuscript, which was commented on, edited, and approved by all authors.

---

**Figure S1.**
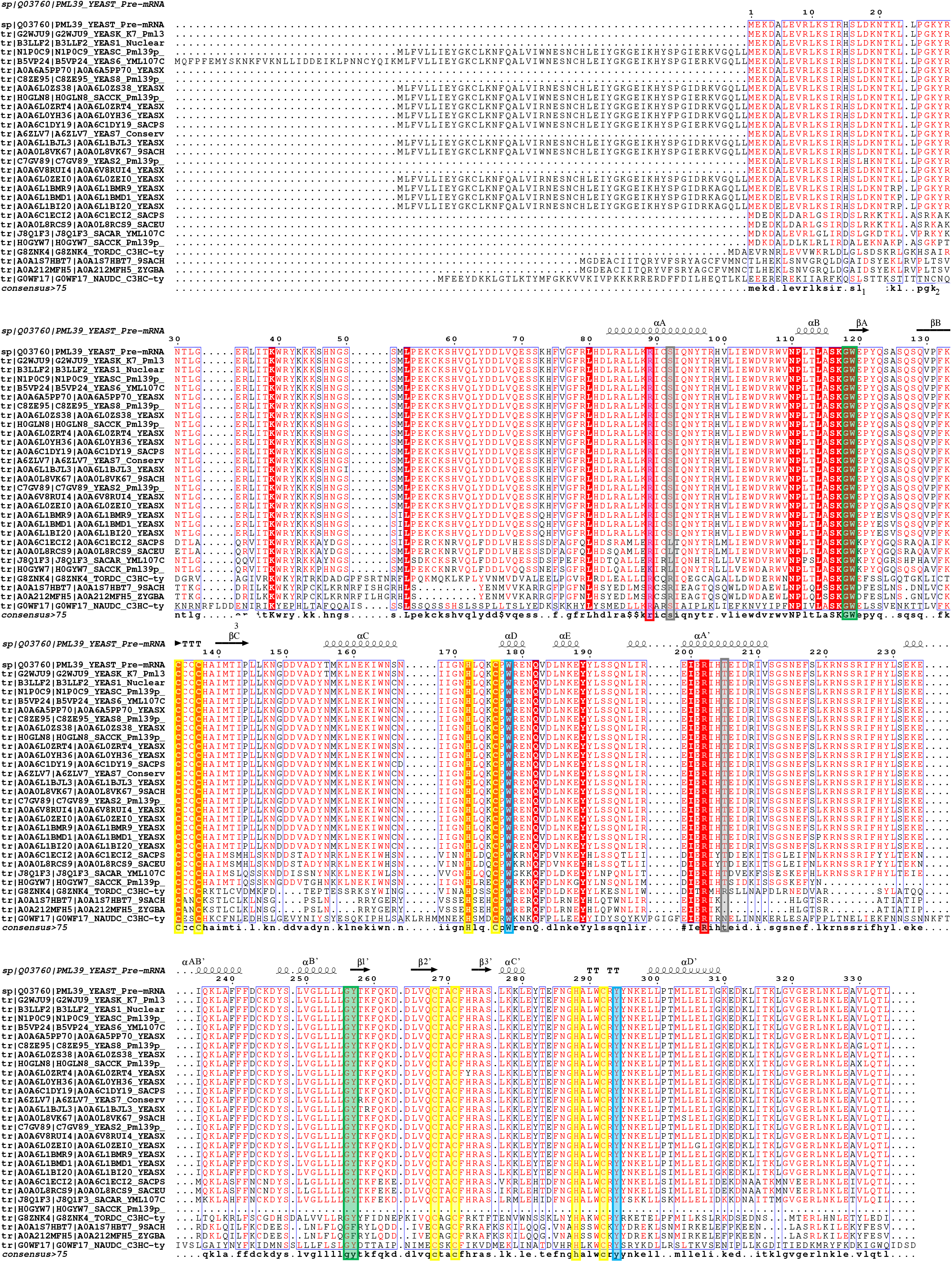
Alignment of *Saccharomycetaceae* Pml39 sequences. Strict identity is indicated by a red box with a white character and similarity is indicated by a red character. Key residues are highlighted by colored boxes according to Fig. 2. Uniprot (https://www.uniprot.org/) accession codes: Q03760|PML39_YEAST Pre-mRNA leakage protein 39 *S. cerevisiae* (strain ATCC 204508 / S288c), Uniprot: G2WJU9|YEASK K7_Pml39p *S. cerevisiae* (strain Kyokai no. 7 / NBRC 101557), Uniprot: B3LLF2|YEAS1 Nuclear pore-associated protein *S. cerevisiae* (strain RM11-1a) Uniprot: C8ZE95|YEAS8 Pml39p *S. cerevisiae* (strain Lalvin EC1118 / Prise de mousse), Uniprot: B5VP24|YEAS6 YML107Cp-like protein *S. cerevisiae* (strain AWRI1631), Uniprot: N1P0C9|Pml39p *S. cerevisiae* (strain CEN.PK113-7D), Uniprot: H0GLN8|SACCK Pml39p *S. cerevisiae* x Saccharomyces kudriavzevii (strain VIN7), Uniprot: A0A0L8VK67|9SACH PML39p Protein required for nuclear retention of unspliced pre-mRNAs *S. sp. ‘boulardii’*, Uniprot: A6ZLV7|YEAS7 Conserved protein *S. cerevisiae* (strain YJM789), Uniprot: C7GV89| YEAS2 Pml39p *S. cerevisiae* (strain JAY291), Uniprot: A0A0L8RCS9| SACEU PML39-like *protein S. eubayanus*, Uniprot: J8Q1F3|SACAR YML107C *S. arboricola* (strain H-6 / AS 2.3317 / CBS 10644), Uniprot: H0GYW7| SACCK Pml39p *S. cerevisiae* x Saccharomyces kudriavzevii (strain VIN7), Uniprot: G8ZNK4| TORDC Uncharacterized protein *Torulaspora delbrueckii* (strain ATCC 10662 / CBS 1146 / NBRC 0425 / NCYC 2629 / NRRL Y-866), Uniprot: S6E752| ZYGB2 ZYBA0S04-07140g1_1 *Zygosaccharomyces bailii* (strain CLIB 213 / ATCC 58445 / CBS 680 / CCRC 21525 / NBRC 1098 / NCYC 1416 / NRRL Y-2227), Uniprot: A0A1S7HBT7| 9SACH PML39 (YML107C) OS=*Zygosaccharomyces* parabailii, Uniprot: A0A212MFH5| ZYGBA Uncharacterized protein *Zygosaccharomyces* bailii, Uniprot: G0WF17| NAUDC Uncharacterized protein *Naumovozyma* dairenensis (strain ATCC 10597 / BCRC 20456 / CBS 421 / NBRC 0211 / NRRL Y-12639). 75% consensus sequence is shown on the bottom.

**Figure S2.**
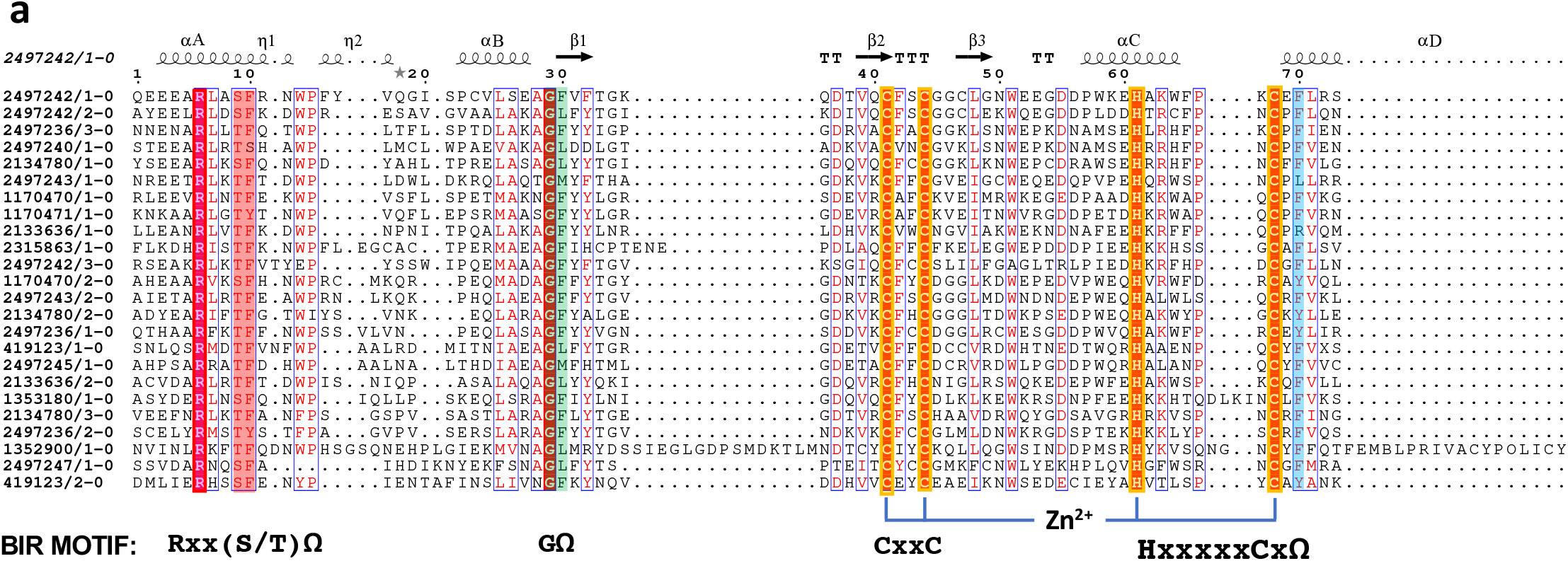
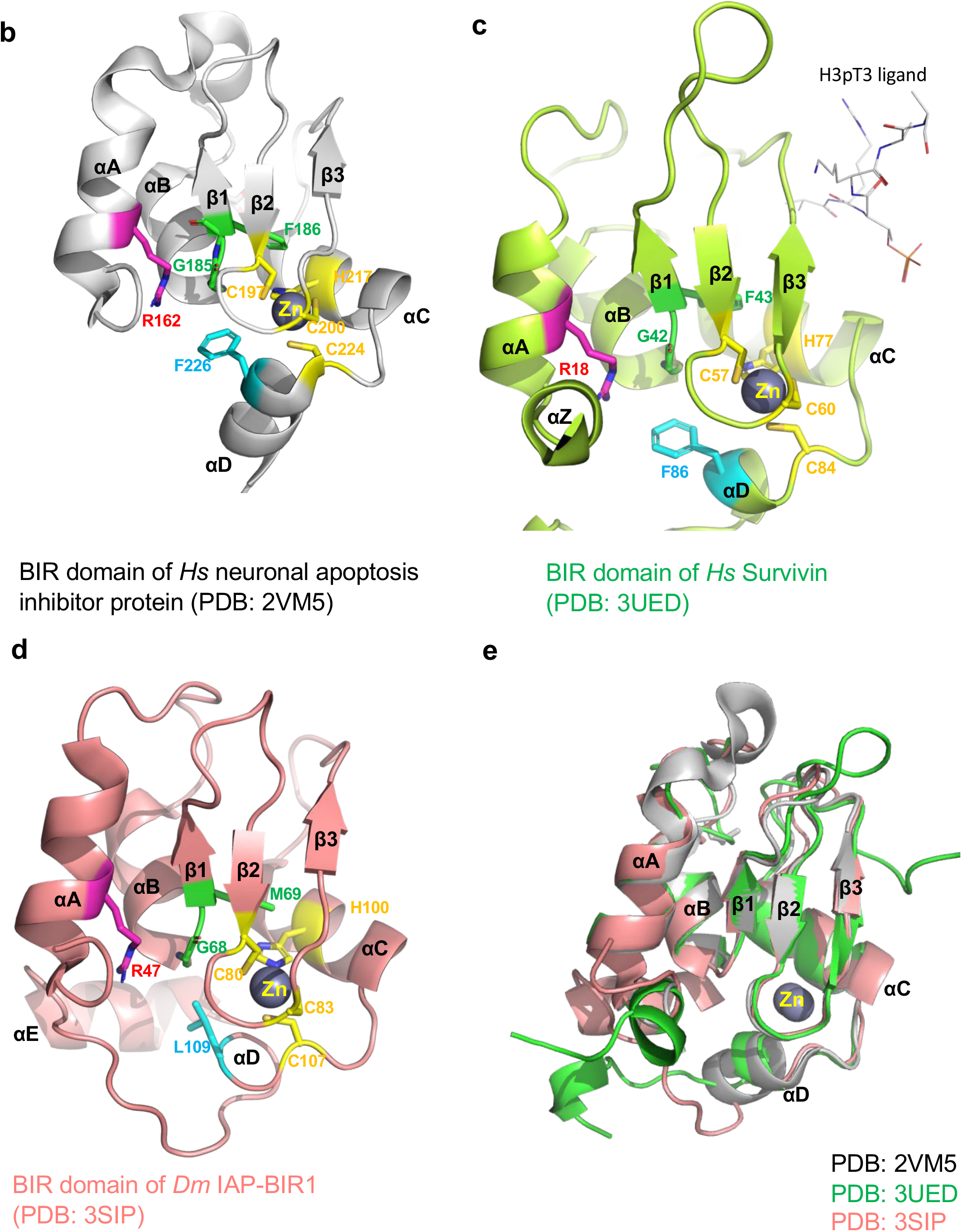
Sequence and structure alignment of canonical BIR domains. **a)** For each BIR domain, the sequences are referred to by their NCBI databank ID (https://www.ncbi.nlm.nih.gov/protein) and the number of the BIR domain if multiple BIR domains are present. Key residues are colored according to Fig. 2. The secondary structure of BIR1 domain of neuronal apoptosis inhibitory protein (PDB: 2VM5) is shown on top. The protein names and Uniprot (U) or Genbank (GB) accession codes are as follows: neuronal apoptosis inhibitory protein birl (U: Q13075), neuronal apoptosis inhibitory protein bir2 (U: Q13075), Baculoviral IAP repeat-containing protein 3 BIR3 (U: Q13489.2) Inhibitor of apoptosis protein BIR1 (U: Q90660.1), E3 ubiquitin-protein ligase XIAP BIR1 (U: P98170.2), Apoptosis 1 inhibitor (Inhibitor of apoptosis 1) (dIAP1) (Thread protein) BIR1 (U: Q24306), Apoptosis inhibitor IAP BIR1 (U: P41436.1), E3 ubiquitin-protein ligase IAP-3 BIR1 (U: P41437.1), Death-associated inhibitor of apoptosis 2 BIR1 (U: Q24307.3), apoptosis inhibitor survivin BIR2 (GB: AAC51660.1), neuronal apoptosis inhibitory protein bir3 (U: Q13075), Apoptosis inhibitor IAP BIR2 (U: P41436.1), Apoptosis 1 inhibitor (Inhibitor of apoptosis 1) (dIAP1) (Thread protein) BIR2 (U: Q24306), E3 ubiquitin-protein ligase XIAP BIR2 (U: P98170.2), Baculoviral IAP repeat-containing protein 3 BIR1 (U: Q13489.2), Apoptosis inhibitor 1 BIR1 (U: P41435.1), Death-associated inhibitor of apoptosis 2 BIR2 (U: Q24307.3), Apoptosis inhibitor 193R BIR1 (U: P47732.2), E3 ubiquitin-protein ligase XIAP BIR3 (U: P98170.2), Baculoviral IAP repeat-containing protein 3 BIR2 (U: Q13489.2), *S. cerevisiae* BIR1 BIR1 (U: P47134.1), IAP-like protein p27 BIR1 (U: P68763.1), Apoptosis inhibitor 1 BIR2 (U: P41435.1). **b)** Crystal structure of the canonical BIR1 domain of neuronal apoptosis inhibitory protein (PDB: 2VM5). **c)** Crystal structure of BIR domain of *D. melanogaster* IAP-BIR1 (PDB: 3SIP). **d)** Crystal structure of BIR domain of Hs Survivin (PDB: 3UED). **e)** Superimposition of these structures.

**Figure S3.**
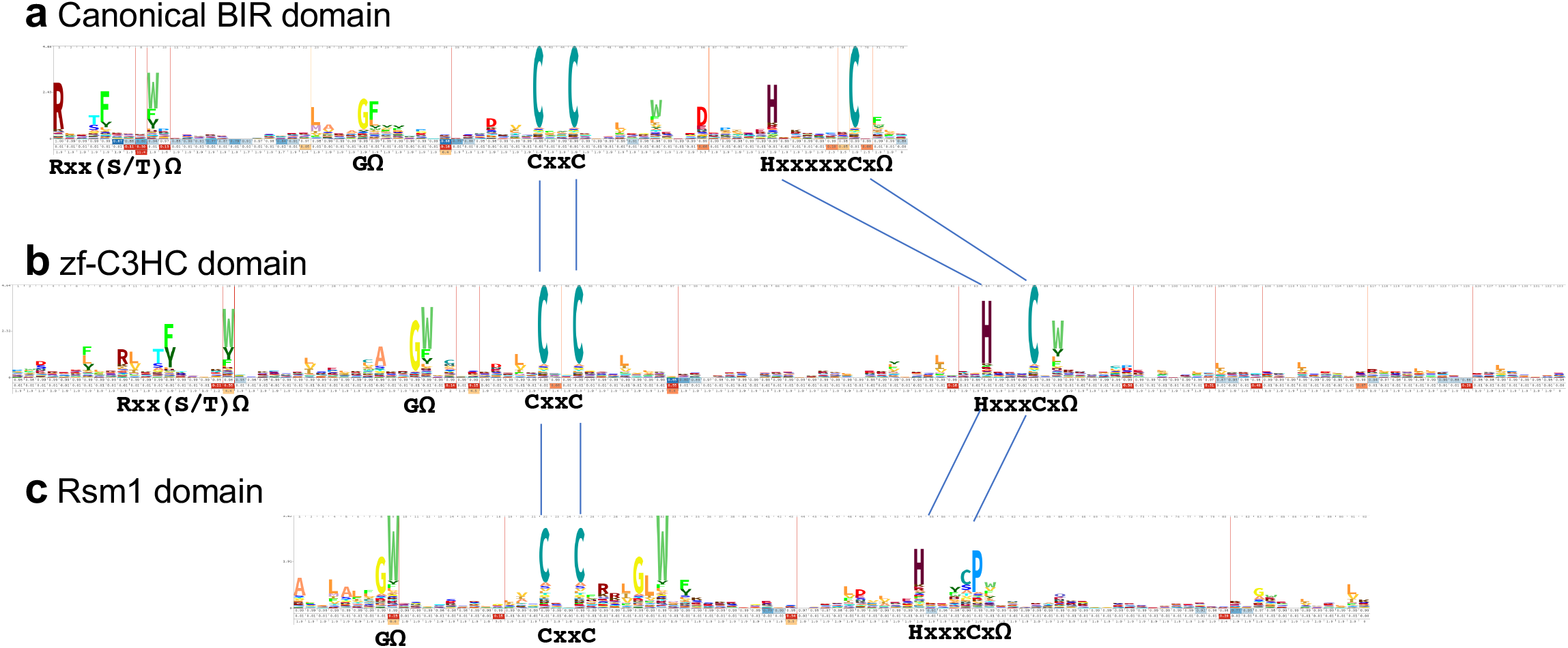
Consensus sequences of BIR-like domains. Hidden Markov model of the **a)** canonical BIR domain, **b)** zf-C3HC domain, and **c)** Rsm1 domain.

**Figure S4.**
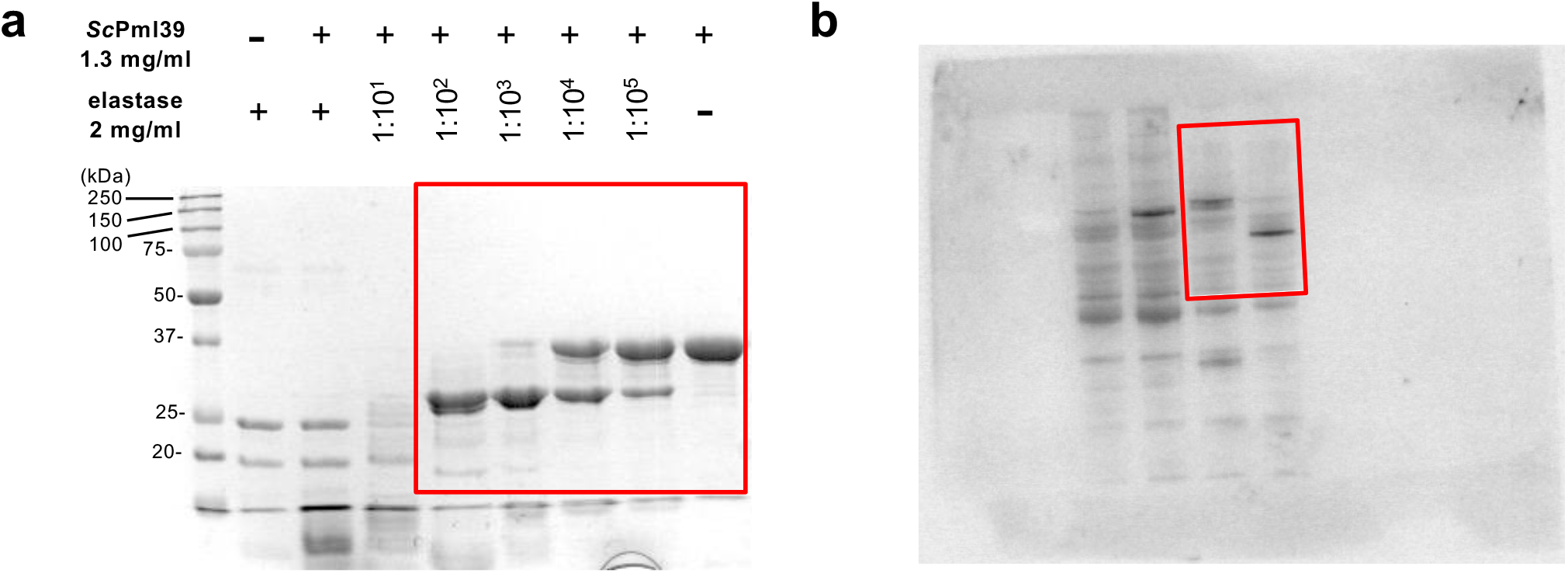
Full gel and immunoblot related to Figure 1. **a)** Full gel of cropped gel displayed in Fig. 1a indicated by the red box. **b)** Full immunoblot of cropped blot displayed in Fig. 1b indicated by the red box.

**Figure S5.**
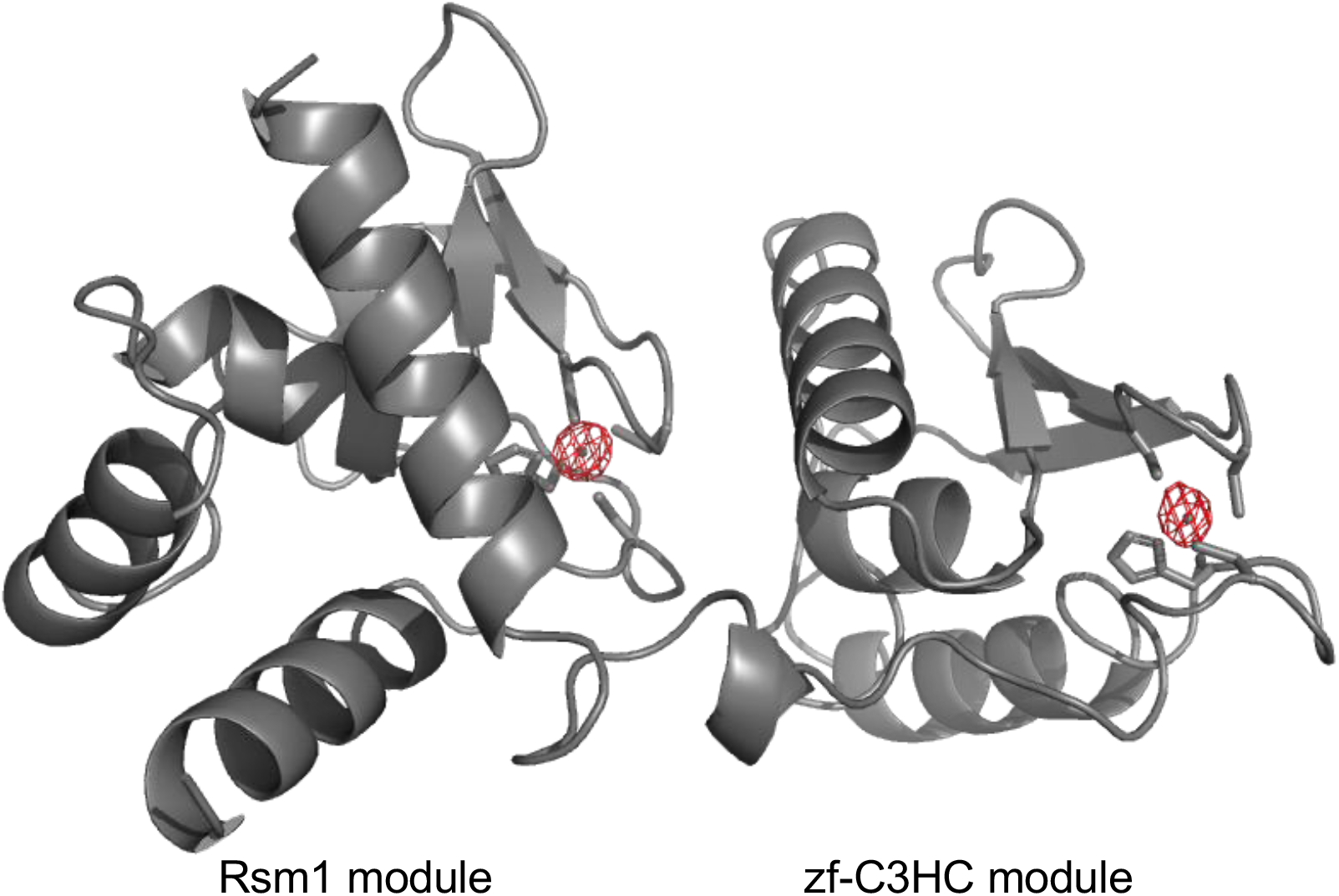
Anomalous Zn difference map of *Sc*Pml39_77-317_ contoured in a red mesh at 10σ.

**Figure S6.**
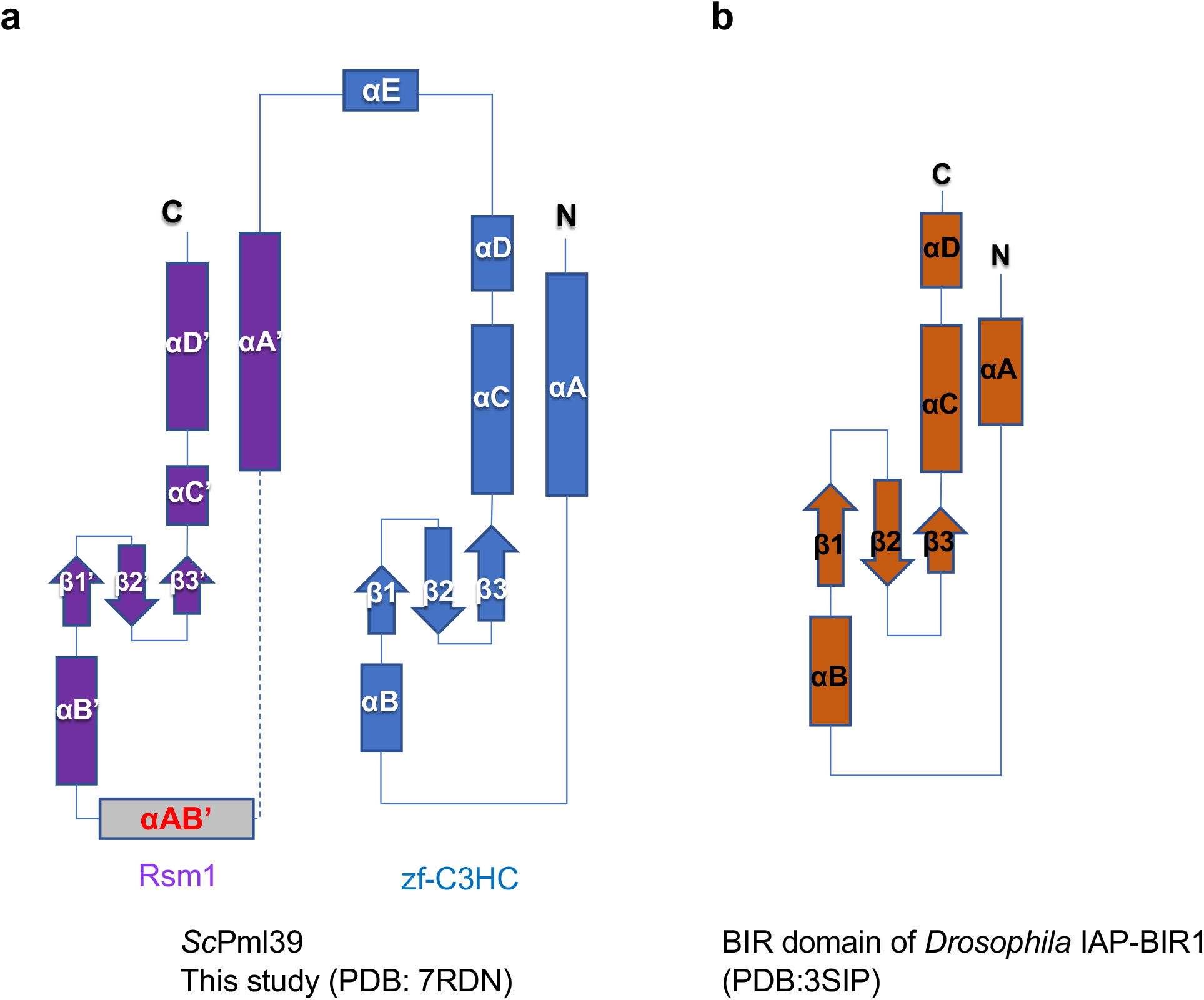
Topology of *Sc*Pml39 (PDB: 7RDN) and BIR domain of human Survivin (3UED).

**Figure S7.**
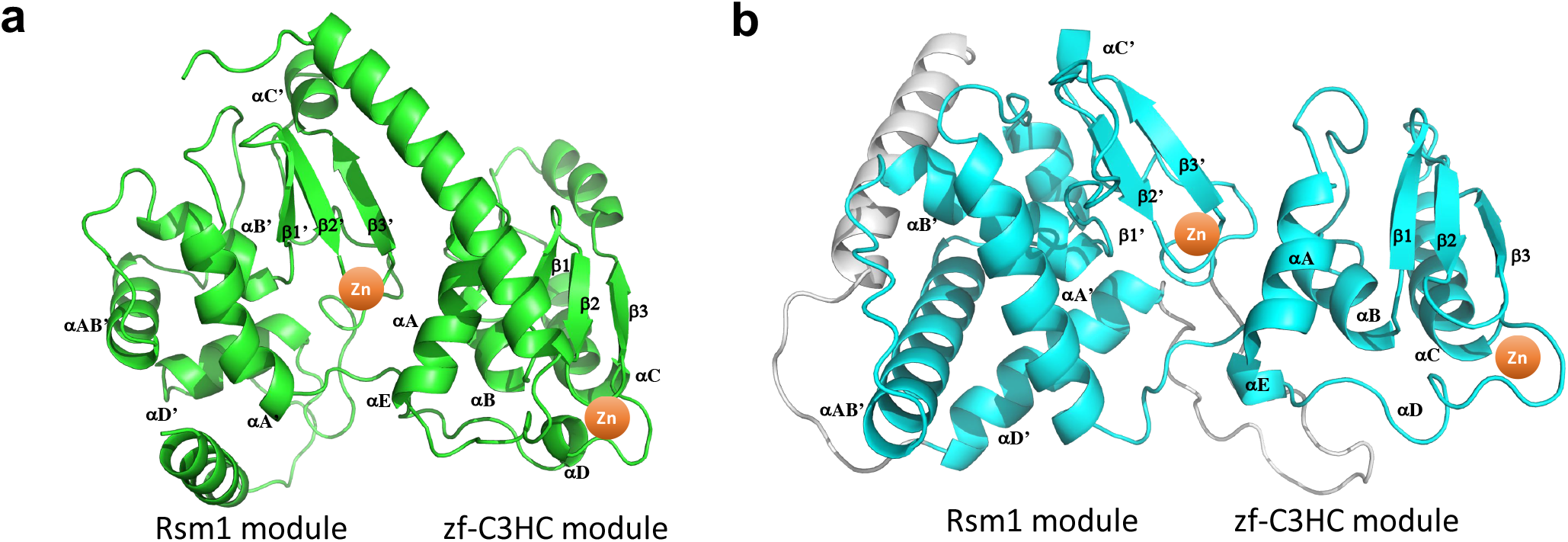
AlphaFold2 models of. **a)** *Sp*Rsm1 (UniProt accession code: O94506, https://alphafold.ebi.ac.uk/entry/O94506) and **b)** *Hs*NIPA/ZC3HC1, splice variant 1 (Q86WB0, https://alphafold.ebi.ac.uk/entry/Q86WB0). Long *Hs*NIPA/ ZC3HC1 loops are omitted for clarity.

**Figure S8.**
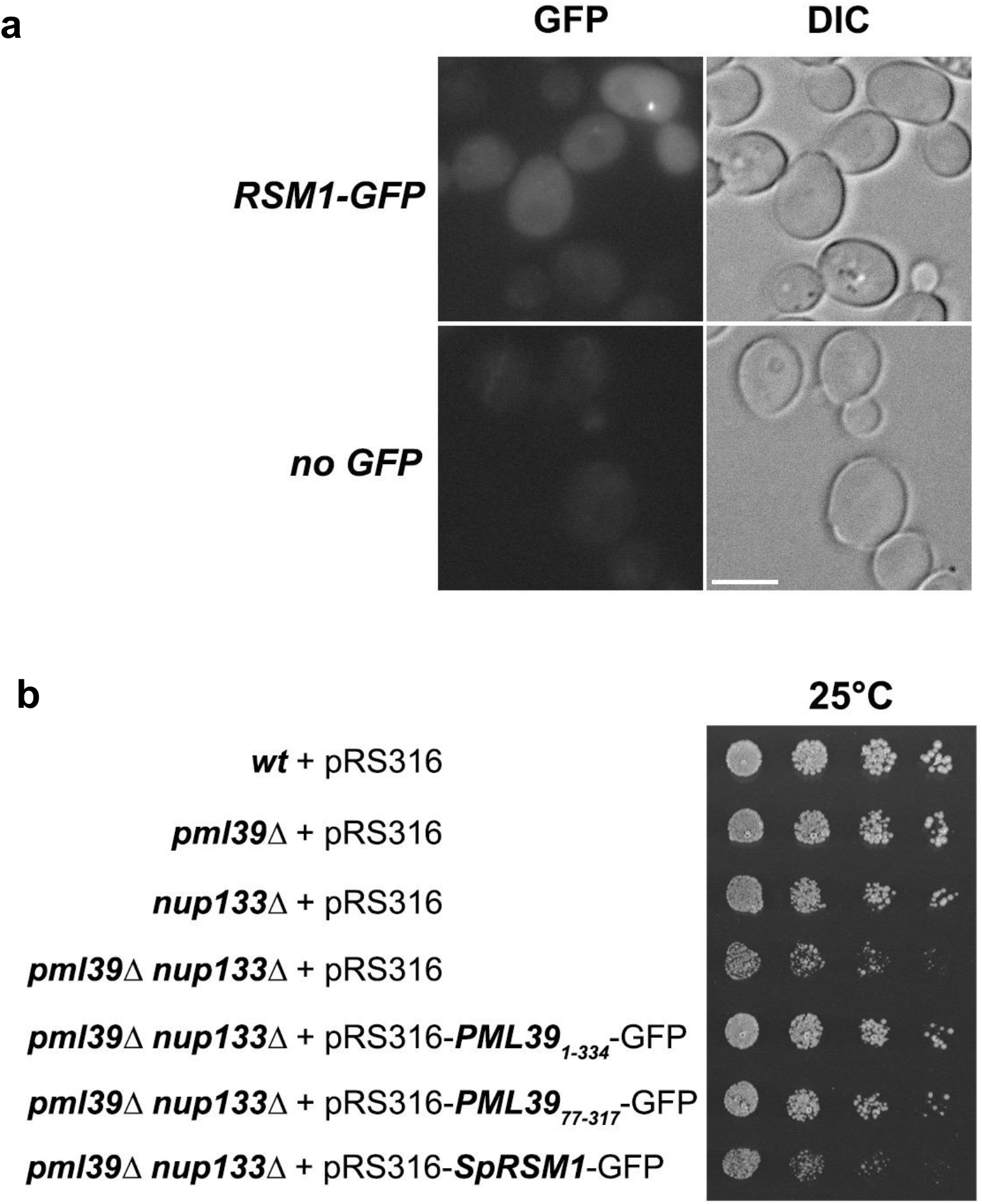
Expression of *Schizosaccharomyces pombe* Rsm1 in *S. cerevisiae*. **a)** *Sp*Rsm1 does not localize at the nuclear periphery when expressed in budding yeast. *S. cerevisiae pml39*Δ cells carrying an empty vector (*“no GFP”*) or expressing GFP-tagged Rsm1 under the control of the *PML39* promoter (top panel) were imaged as in Fig. 1. Single plane images are shown for the GFP and DIC (differential interference contrast) channels. Scale bar, 5 μm. **b)** *Sp*Rsm1 does not complement the *pml39Δ nup133*Δ synthetic interaction in the growth assay. Cells of the indicated genotypes were spotted as serial dilutions on SC medium and grown for 3 days at 25°C (same spotting as in Fig. 1d).

**Figure S9.**
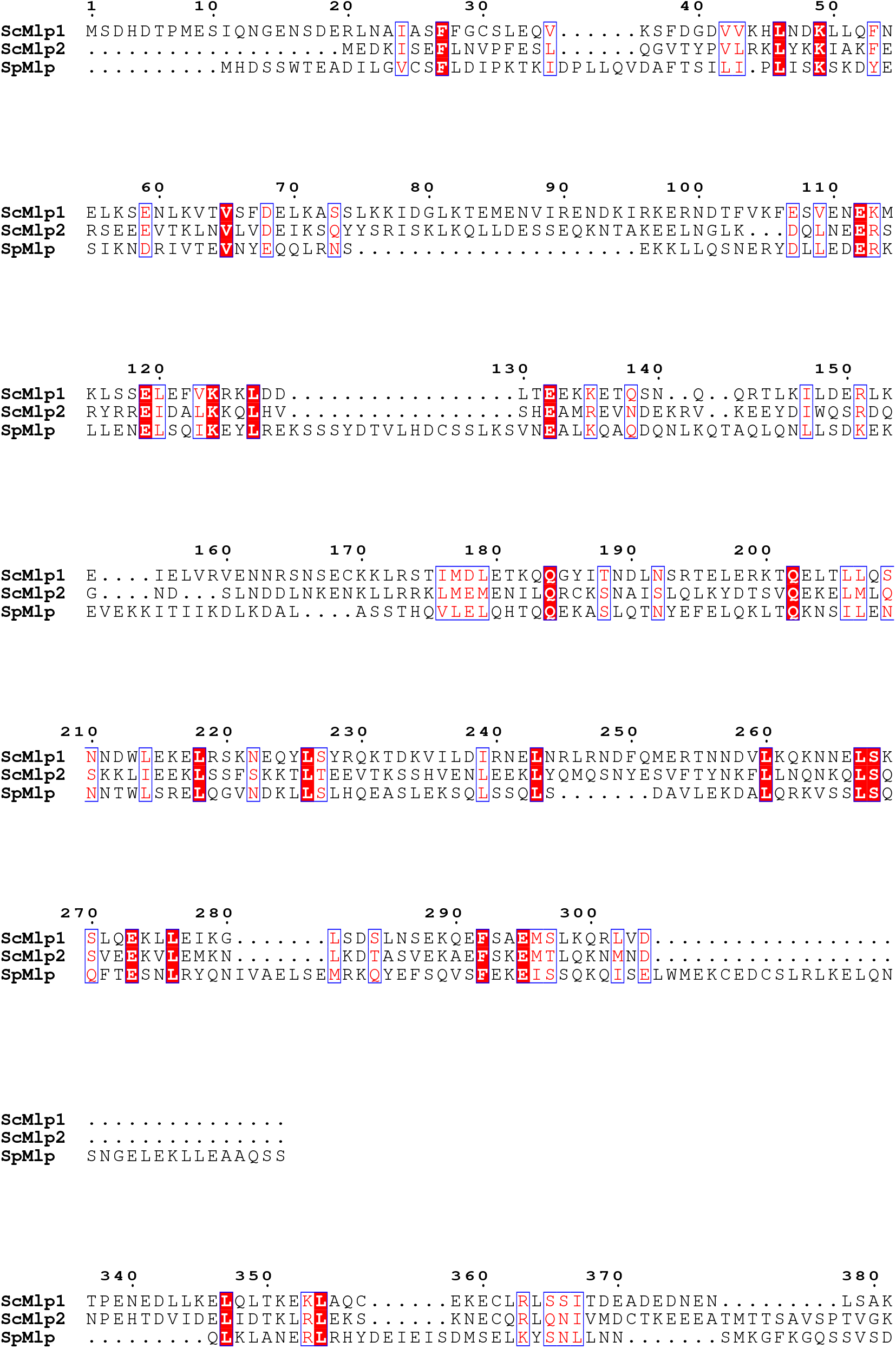
Sequence alignment among N-terminal *S. cerevisiae* Mlp1, *S. cerevisiae* Mlp2, and *S. pombe* nup211 (*Sp*Mlp) regions. *Sc*Pml39 interacts with dimeric *Sc*Mlp1_1-350_ *in vitro*. UniProt accession codes: Q02455 (*Sc*Mlp1), P40457 (*S*cMlp2), and O74424 (*Sp*Nup211, referred to as SpMlp here).

**Table S1.**
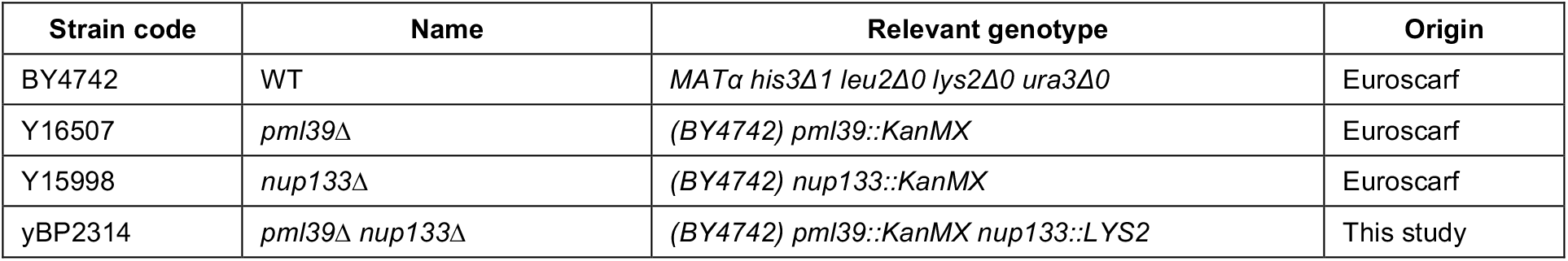
Yeast Strains used in this study.

## References

1. Wente, S. R. & Rout, M. P. The nuclear pore complex and nuclear transport. Cold Spring Harb. Perspect. Biol. 2, a000562 (2010).

2. Hoelz, A., Debler, E. W. & Blobel, G. The structure of the nuclear pore complex. Annu. Rev. Biochem. 80, 613–43 (2011).

3. Schwartz, T. U. The Structure Inventory of the Nuclear Pore Complex. J. Mol. Biol. 428, 1986–2000 (2016).

4. Lin, D. H. & Hoelz, A. The structure of the nuclear pore complex (An Update). Annu. Rev. Biochem. 88, 725–783 (2019).

5. Blobel, G. Gene gating: a hypothesis. Proc. Natl. Acad. Sci. 82, 8527–8529 (1985).

6. Akhtar, A. & Gasser, S. M. The nuclear envelope and transcriptional control. Nat. Rev. Genet. 8, 507–517 (2007).

7. Köhler, A. & Hurt, E. Gene Regulation by Nucleoporins and Links to Cancer. Mol. Cell 38, 6–15 (2010).

8. Sood, V. & Brickner, J. H. Nuclear pore interactions with the genome. Curr. Opin. Genet. Dev. 25, 43–49 (2014).

9. Ptak, C. & Wozniak, R. W. Nucleoporins and chromatin metabolism. Curr. Opin. Cell Biol. 40, 153–160 (2016).

10. Pascual-Garcia, P. & Capelson, M. The nuclear pore complex and the genome: organizing and regulatory principles. Curr. Opin. Genet. Dev. 67, 142–150 (2021).

11. Bonnet, A. & Palancade, B. Regulation of mRNA Trafficking by Nuclear Pore Complexes. Genes (Basel). 5, 767–791 (2014).

12. Buchwalter, A., Kaneshiro, J. M. & Hetzer, M. W. Coaching from the sidelines: the nuclear periphery in genome regulation. Nat. Rev. Genet. 20, 39–50 (2019).

13. Schmid, M. & Jensen, T. H. Transcription-associated quality control of mRNP. Biochim. Biophys. Acta 1829, 158–168 (2013).

14. Wolin, S. L. & Maquat, L. E. Cellular RNA surveillance in health and disease. Science 366, 822–827 (2019).

15. Green, D. M., Johnson, C. P., Hagan, H. & Corbett, A. H. The C-terminal domain of myosin-like protein 1 (Mlp1p) is a docking site for heterogeneous nuclear ribonucleoproteins that are required for mRNA export. Proc. Natl. Acad. Sci. 100, 1010–1015 (2003).

16. Vinciguerra, P., Iglesias, N., Camblong, J., Zenklusen, D. & Stutz, F. Perinuclear Mlp proteins downregulate gene expression in response to a defect in mRNA export. EMBO J. 24, 813–823 (2005).

17. Stewart, M. Nuclear export of mRNA. Trends Biochem. Sci. 35, 609–617 (2010).

18. Niepel, M. et al. The nuclear basket proteins Mlp1p and Mlp2p are part of a dynamic interactome including Esc1p and the proteasome. Mol. Biol. Cell 24, 3920–3938 (2013).

19. Saroufim, M. A. et al. The nuclear basket mediates perinuclear mRNA scanning in budding yeast. J. Cell Biol. 211, 1131–1140 (2015).

20. Hector, R. E. et al. Dual requirement for yeast hnRNP Nab2p in mRNA poly(A) tail length control and nuclear export. EMBO J. 21, 1800–1810 (2002).

21. Grant, R. P. et al. Structure of the N-Terminal Mlp1-Binding Domain of the Saccharomyces cerevisiae mRNA-Binding Protein, Nab2. J. Mol. Biol. 376, 1048–1059 (2008).

22. Brockmann, C. et al. Structural basis for polyadenosine-RNA binding by Nab2 Zn fingers and its function in mRNA nuclear export. Structure 20, 1007–1018 (2012).

23. Galy, V. et al. Nuclear Retention of Unspliced mRNAs in Yeast Is Mediated by Perinuclear Mlp1. Cell 116, 63–73 (2004).

24. Lo, C. W. et al. Inhibition of splicing and nuclear retention of pre-mRNA by spliceostatin A in fission yeast. Biochem. Biophys. Res. Commun. 364, 573–577 (2007).

25. Coyle, J. H., Bor, Y. C., Rekosh, D. & Hammarskjold, M. L. The Tpr protein regulates export of mRNAs with retained introns that traffic through the Nxf1 pathway. RNA 17, 1344–1356 (2011).

26. Rajanala, K. & Nandicoori, V. K. Localization of nucleoporin Tpr to the nuclear pore complex is essential for Tpr mediated regulation of the export of unspliced RNA. PLoS One 7, e29921 (2012).

27. Palancade, B., Zuccolo, M., Loeillet, S., Nicolas, A. & Doye, V. Pml39, a Novel Protein of the Nuclear Periphery Required for Nuclear Retention of Improper Messenger Ribonucleoparticles. Mol. Biol. Cell 16, 5258–5268 (2005).

28. Siniossoglou, S. et al. A novel complex of nucleoporins, which includes Sec13p and a Sec13p homolog, is essential for normal nuclear pores. Cell 84, 265–275 (1996).

29. Debler, E. W., Hsia, K.-C., Nagy, V., Seo, H.-S. & Hoelz, A. Characterization of the membrane-coating Nup84 complex: paradigm for the nuclear pore complex structure. Nucleus 1, 150–7 (2010).

30. Fasken, M. B., Corbett, A. H. & Stewart, M. Structure-function relationships in the Nab2 polyadenosine-RNA binding Zn finger protein family. Protein Sci. 28, 513–523 (2019).

31. Monzon, V., Paysan-Lafosse, T., Wood, V. & Bateman, A. Reciprocal Best Structure Hits: Using AlphaFold models to discover distant homologues. bioRxiv 2022.07.04.498216 (2022). doi:10.1101/2022.07.04.498216

32. Waterhouse, A. et al. SWISS-MODEL: homology modelling of protein structures and complexes. Nucleic Acids Res. 46, W296–W303 (2018).

33. Kelley, L. A., Mezulis, S., Yates, C. M., Wass, M. N. & Sternberg, M. J. E. The Phyre2 web portal for protein modeling, prediction and analysis. Nat. Protoc. 10, 845–58 (2015).

34. Takahashi, R. et al. A single BIR domain of XIAP sufficient for inhibiting caspases. J. Biol. Chem. 273, 7787–90 (1998).

35. Deveraux, Q. L., Takahashi, R., Salvesen, G. S. & Reed, J. C. X-linked IAP is a direct inhibitor of cell-death proteases. Nature 388, 300–304 (1997).

36. Cossu, F., Milani, M., Mastrangelo, E. & Lecis, D. Targeting the BIR Domains of Inhibitor of Apoptosis (IAP) Proteins in Cancer Treatment. Comput. Struct. Biotechnol. J. 17, 142–150 (2019).

37. Verhagen, A. M., Coulson, E. J. & Vaux, D. L. Inhibitor of apoptosis proteins and their relatives: IAPs and other BIRPs. Genome Biol. 2, REVIEWS3009 (2001).

38. Ouyang, T. et al. Identification and characterization of a nuclear interacting partner of anaplastic lymphoma kinase (NIPA). J. Biol. Chem. 278, 30028–30036 (2003).

39. Bourhis, E., Hymowitz, S. G. & Cochran, A. G. The mitotic regulator Survivin binds as a monomer to its functional interactor Borealin. J. Biol. Chem. 282, 35018–23 (2007).

40. Du, J., Kelly, A. E., Funabiki, H. & Patel, D. J. Structural basis for recognition of H3T3ph and Smac/DIABLO N-terminal peptides by human Survivin. Structure 20, 185–95 (2012).

41. Niedzialkowska, E. et al. Molecular basis for phosphospecific recognition of histone H3 tails by Survivin paralogues at inner centromeres. Mol. Biol. Cell 23, 1457–66 (2012).

42. Mace, P. D., Shirley, S. & Day, C. L. Assembling the building blocks: structure and function of inhibitor of apoptosis proteins. Cell Death Differ. 17, 46–53 (2010).

43. Luque, L. E., Grape, K. P. & Junker, M. A highly conserved arginine is critical for the functional folding of inhibitor of apoptosis (IAP) BIR domains. Biochemistry 41, 13663–13671 (2002).

44. Mistry, J. et al. Pfam: The protein families database in 2021. Nucleic Acids Res. 49, D412–D419 (2021).

45. Wheeler, T. J., Clements, J. & Finn, R. D. Skylign: a tool for creating informative, interactive logos representing sequence alignments and profile hidden Markov models. BMC Bioinformatics 15, 7 (2014).

46. Laity, J. H., Lee, B. M. & Wright, P. E. Zinc finger proteins: New insights into structural and functional diversity. Curr. Opin. Struct. Biol. 11, 39–46 (2001).

47. Vucic, D., Kaiser, W. J. & Miller, L. K. A mutational analysis of the baculovirus inhibitor of apoptosis Op-IAP. J. Biol. Chem. 273, 33915–33921 (1998).

48. Birnbaum, M. J., Clem, R. J. & Miller, L. K. An apoptosis-inhibiting gene from a nuclear polyhedrosis virus encoding a polypeptide with Cys/His sequence motifs. J. Virol. 68, 2521–2528 (1994).

49. Deveraux, Q. L. & Reed, J. C. IAP family proteins--suppressors of apoptosis. Genes Dev. 13, 239–52 (1999).

50. Yoon, J. H. Schizosaccharomyces pombe rsm1 genetically interacts with spmex67, which is involved in mRNA export. J. Microbiol. 42, 32–36 (2004).

51. Moon, D. G. R. M., Park, Y. S., Kim, C. Y. & Yoon, J. H. Isolation of synthetic lethal mutations in the rsm1-null mutant of fission yeast. J. Microbiol. 48, 701–705 (2010).

52. Bassermann, F. et al. NIPA defines an SCF-type mammalian E3 ligase that regulates mitotic entry. Cell 122, 45–57 (2005).

53. Kokoszynska, K., Rychlewski, L. & Wyrwicz, L. S. The mitotic entry regulator NIPA is a prototypic BIR domain protein. Cell Cycle 7, 2073–2075 (2008).

54. Jumper, J. et al. Highly accurate protein structure prediction with AlphaFold. Nature 596, 583–589 (2021).

55. Bonnet, A., Bretes, H. & Palancade, B. Nuclear pore components affect distinct stages of intron-containing gene expression. Nucleic Acids Res. 43, 4249–4261 (2015).

56. Gunkel, P., Iino, H., Krull, S. & Cordes, V. C. ZC3HC1 Is a Novel Inherent Component of the Nuclear Basket, Resident in a State of Reciprocal Dependence with TPR. Cells 10, 1937 (2021).

57. Gunkel, P. & Cordes, V. C. ZC3HC1 is a structural element of the nuclear basket effecting interlinkage of TPR polypeptides. Mol. Biol. Cell 33, ar82 (2022).

58. Peri, S., Steen, H. & Pandey, A. GPMAW--a software tool for analyzing proteins and peptides. Trends Biochem. Sci. 26, 687–689 (2001).

59. Otwinowski, Z. & Minor, W. Macromolecular Crystallography Part A. Methods Enzymol. 276, 307–326 (1997).

60. Liebschner, D. et al. Macromolecular structure determination using X-rays, neutrons and electrons: Recent developments in Phenix. Acta Crystallogr. Sect. D Struct. Biol. 75, 861–877 (2019).

61. Jones, T. A., Zou, J. -Y., Cowan, S. W. & Kjeldgaard, M. Improved methods for building protein models in electron density maps and the location of errors in these models. Acta Crystallogr. Sect. A 47, 110–119 (1991).

62. Emsley, P., Lohkamp, B., Scott, W. G. & Cowtan, K. Features and development of Coot. Acta Crystallogr. Sect. D Biol. Crystallogr. 66, 486–501 (2010).

63. Williams, C. J. et al. MolProbity: More and better reference data for improved all-atom structure validation. Protein Sci. 27, 293–315 (2018).

64. Baker, N. A., Sept, D., Joseph, S., Holst, M. J. & McCammon, J. A. Electrostatics of nanosystems: Application to microtubules and the ribosome. Proc. Natl. Acad. Sci. 98, 10037–10041 (2001).

65. Longtine, M. S. et al. Additional modules for versatile and economical PCR-based gene deletion and modification in Saccharomyces cerevisiae. Yeast 14, 953–961 (1998).

66. Lautier, O. et al. Co-translational assembly and localized translation of nucleoporins in nuclear pore complex biogenesis. Mol. Cell 81, 2417–2427.e5 (2021).

